# Exploring the intricacies and pitfalls of the ATN framework: An assessment across cohorts and thresholding methodologies

**DOI:** 10.1101/2022.12.06.519269

**Authors:** Yasamin Salimi, Daniel Domingo-Fernández, Martin Hofmann-Apitius, Colin Birkenbihl, the Alzheimer’s Disease Neuroimaging Initiative, the Japanese Alzheimer’s Disease Neuroimaging Initiative, the Alzheimer’s Disease Repository Without Borders Investigators, the European Prevention of Alzheimer’s Disease (EPAD) Consortium

## Abstract

The amyloid/tau/neurodegeneration (ATN) framework has redefined Alzheimer’s disease (AD) toward a primarily biological entity. While it has found wide application in AD research, it was so far typically applied to single cohort studies using distinct data-driven thresholding methods. This poses the question of how concordant thresholds obtained using distinct methods are within the same dataset as well as whether thresholds derived by the same technique are interchangeable across cohorts. Given potential differences in cohort data-derived thresholds, it remains unclear whether individuals of one cohort are actually comparable with regard to their exhibited disease patterns to individuals of another cohort, even when they are assigned to the same ATN profile. If such comparability is not evident, the generalizability of results obtained using the ATN framework is at question. In this work, we evaluated the impact of selecting specific thresholding methods on ATN profiles by applying five commonly-used methodologies across eleven AD cohort studies. Our findings revealed high variability among the obtained thresholds, both across methods and datasets, linking the choice of thresholding method directly to the type I and type II error rate of ATN profiling. Moreover, we assessed the generalizability of primarily Magnetic Resonance Imaging (MRI) derived biological patterns discovered within ATN profiles by simultaneously clustering participants of different cohorts who were assigned to the same ATN profile. In only two out of seven investigated ATN profiles, we observed a significant association between individuals’ assigned clusters and cohort origin for thresholds defined using Gaussian Mixture Models, while no significant associations were found for K-means-derived thresholds. Consequently, in the majority of profiles, biological signals governed the clustering rather than systematic cohort differences resulting from distinct biomarker thresholds. Our work revealed that: 1) the thresholding method selection is a decision of statistical relevance that will inevitably bias the resulting profiling, 2) obtained thresholds are most likely not directly interchangeably across two independent cohorts, and 3) MRI-based biological patterns derived from distinctly thresholded ATN profiles can generalize across cohort datasets. Conclusively, in order to appropriately apply the ATN framework as an actionable and robust biological profiling scheme, a comprehensive understanding of the impact of used thresholding methods, their statistical implications, and the validation of achieved results is crucial.

## 1. Background

Alzheimer’s disease (AD) is a progressive condition in which symptoms manifest years after the initial onset of the disease (Jack *et al*. 2018). Over the past decades, AD diagnosis relied predominantly on cognitive assessments (e.g., the Mini-Mental State Examination (MMSE) or Clinical Dementia Rating scale (CDR)), and individuals were commonly assigned to one of the three following diagnostic groups: i) Cognitively Unimpaired (CU), ii) Mild Cognitive Impairment (MCI), and iii) AD. Patients within the cognitively impaired groups, however, exhibit a large degree of heterogeneity with respect to symptoms (Ritchie *et al*. 2001), disease severity, and progression (Birkenbihl *et al*. 2022). One possible reason for this could be that the clinical definition of AD ignores the disease-underlying biological condition of patients such as their state of amyloid burden and neurodegeneration (McKhann *et al*. 2011).

Motivated by these concerns, Jack *et al*. (2018) proposed the β-amyloid deposition (A), pathologic tau (T), and neurodegeneration (N) framework as a potentially unbiased approach for categorizing AD patients according to their biological condition rather than cognitive function. To do so, the ATN framework refers to a specific set of biomarkers (e.g., amyloid positron emission tomography (PET)) to measure whether any of the three factors (i.e., A, T, and N) appear abnormal or not. Based on this categorization scheme, participants are then assigned an abnormal (+), or normal (-) state for each factor, resulting in eight possible biomarker profiles (e.g., A+T+N+, A+T+N-, etc.). To allow for a dichotomization of participants’ biomarker states, a biomarker-specific threshold needs to be defined. To this end, predominantly data-driven techniques have been used in the literature including clustering approaches such as K-means and Gaussian mixture models (GMM), as well as statistical approaches, for example, placing a threshold at a specific quantile of the empirical data.

While over a hundred published studies have employed the ATN framework to categorize participants, the majority of them applied different thresholds, defined using an arbitrary technique, based on single cohorts. However, such data-derived thresholds will inevitably be influenced by the differences in recruitment criteria, data collection procedures, and processing that are found across clinical cohort datasets (Salimi *et al*. 2022) on which the majority of these studies were based. Therefore, to ensure that achieved results do generalize across AD populations, it is essential to evaluate whether disparate methods would result in different thresholds and how far such differences could bias the resulting ATN profiling. Lastly, it remains unclear whether participants of one cohort study would indeed be comparable to another cohort’s participants who were both assigned to the same ATN profile based on respectively different, purely data-determined thresholds.

Recently, multiple ATN-based studies were conducted investigating the selection of biomarkers, the method for defining thresholds (Mattsson-Carlgren *et al*. 2020, Ingala *et al*. 2021), and whether thresholds were interchangeable across two cohorts (Hansson *et al*. 2018). Hansson *et al*. (2018) found that cerebrospinal fluid (CSF) thresholds achieved from the Alzheimer’s Disease Neuroimaging Initiative (ADNI) could be applied to the BioFINDER cohort while adjusting for preanalytical differences. Another recent study by Mattsson-Carlgren *et al*. (2020) inspected potential differences in ATN profiles associated with the choice of biomarkers as well as the method for dichotomization, again using ADNI-derived thresholds on the BioFINDER cohort. They found that there were few differences among the thresholds obtained using different methodologies, except for CSF amyloid-β 1-42 (Aβ1-42) thresholds. Additionally, their findings highlighted that the achieved profiling was altered when different biomarker combinations were used. Finally, a study by Ingala *et al*. (2021) investigated the difference in thresholds reported in the literature and compared those with thresholds that could be obtained from a cohort using different methods. They leveraged the European Prevention of Alzheimer’s Dementia (EPAD) dataset and reported that GMM thresholds aligned with literature-reported values. However, such previous studies focused on a limited number of cohorts, and the question of generalizability, meaning whether disease patterns exhibited by participants assigned to the same ATN profile were comparable across cohorts, was not investigated. In order for the ATN framework to present an actionable, robust, universal, and finally, unbiased profiling scheme based on participants’ biological conditions, a comprehensive understanding of applied thresholding methods, their impact, and the generalizability of achieved results is crucial.

In this work, we identified a set of 11 AD cohorts that contained the CSF biomarkers recommended for applying the ATN framework (i.e., Aβ1-42, phosphorylated tau (pTau), and total tau (tTau)) (Jack *et al*. 2018). Subsequently, we used five well-established methods to define thresholds for each biomarker in each cohort to categorize its participants according to the ATN framework. Following this, we analyzed deviations among the thresholds defined by each method and investigated the impact of such deviations on the underlying profiling. Lastly, we evaluated whether individuals assigned to the same ATN profiles exhibited similar magnetic resonance imaging (MRI) derived patterns across cohorts, despite their different biomarker thresholds. Our analysis highlights how the chosen thresholding method impacts the obtained ATN profiles, resulting in different thresholds and profile assignments. However, despite substantial variation among the data-driven thresholds, we observed comparable biological patterns between independent cohorts’ participants who were assigned to the same ATN profiles.

## 2. Methods

### 2.1 Investigated cohort studies

Using the ADataViewer platform (Salimi *et al*. 2022), we identified a set of 11 cohort studies that measured the CSF biomarkers necessary for ATN profiling their participants **(Table 1)**. We restricted our analysis to CSF biomarkers as this modality was available in an abundant number of studies and, according to the ATN framework, CSF biomarkers alone are considered sufficient for an appropriate ATN profiling (Jack *et al*. 2018).

**Table 1:**
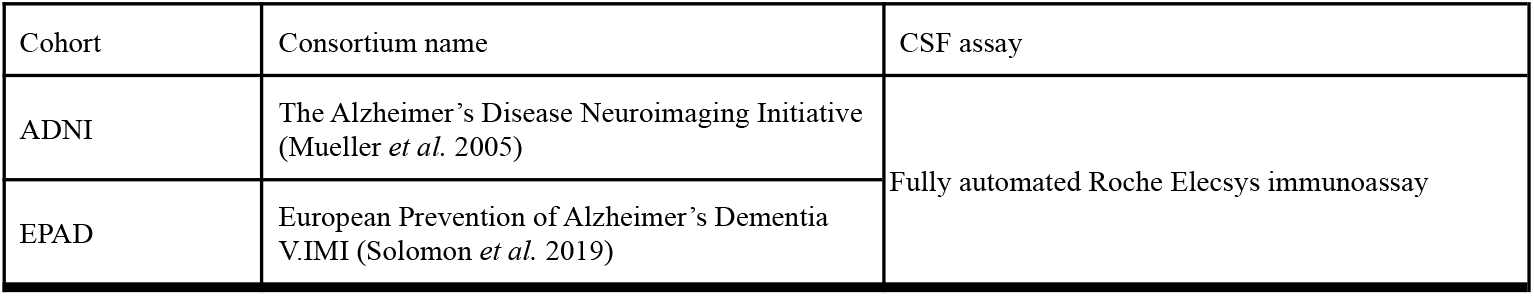
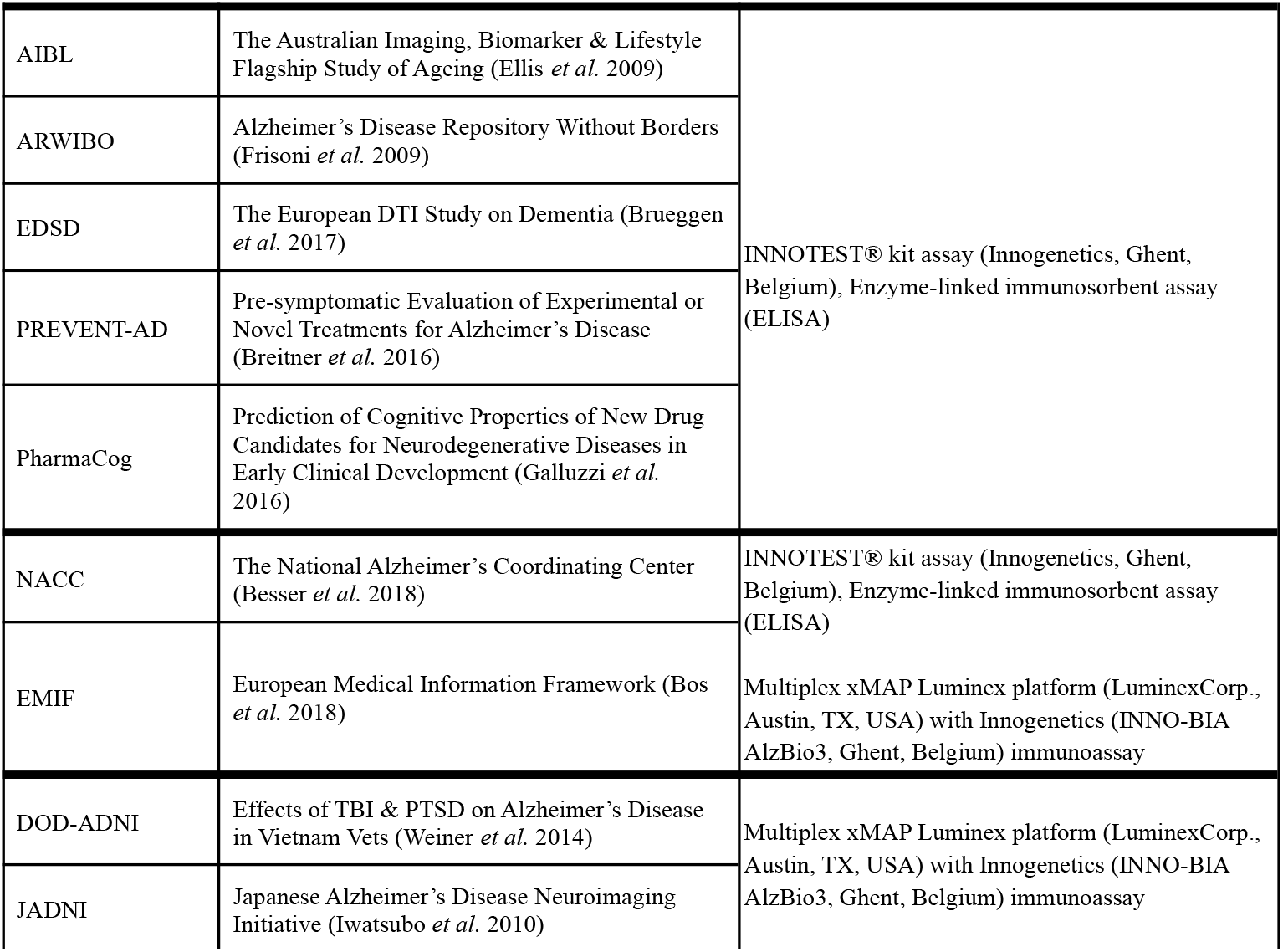
Investigated cohort studies and their respectively employed CSF assays. Cohorts are grouped together if they employed the same assay(s). Note that EMIF and NACC used two distinct assays.

To maximize the number of available participants, we exclusively focused on measurements taken at each cohort studies’ respective baseline. Additionally, we excluded participants with CSF measurements below or above the assay-specific technical limit reported in the study (**Table S1**). The number of remaining participants per cohort as well as summary statistics of key demographic variables is shown in **Table S2**. Lastly, as certain cohorts, specifically NACC and EMIF, contained CSF measurements acquired using different assay methods, we divided their participants into separate groups based on their employed assay **(Table S3)**.

### 2.2 CSF biomarkers

Following the ATN framework recommendations (Jack *et al*. 2018), we selected three CSF biomarkers for profiling participants, namely Aβ1-42, pTau, and tTau indicating A, T, and N, respectively (i.e., N+ indicates that tTau was abnormal while N-indicates that tTau was within a normal range). CSF Aβ1-42 was considered abnormal if the measurement was below a certain threshold value, while CSF pTau and tTau were determined abnormal if the measurement was higher than a given value (McKhann *et al*. 2011, Sperling *et al*. 2011). The assays used to measure these markers in each cohort are presented in **Table 1**. The distribution of each CSF biomarker for cohorts with the same assay is shown in **Fig. S1**.

### 2.3 Thresholding methods

We scanned 116 publications that employed the ATN framework by querying PubMed with [ATN + “Alzheimer’s disease”] with the goal of identifying the most commonly used methods for obtaining biomarker thresholds **(Table S4)**. Our survey revealed the following five data-driven methods as most popular: 1) GMM, 2) K-means clustering, 3) tertile analysis, 4) receiver operating characteristic (ROC) analysis, and 5) mean ±2 standard deviation (SD). Next, we applied these methods to each cohort individually to define a threshold for each of the three biomarkers, resulting in three distinct thresholds per cohort per method. As the interoperability of assays is subject of an ongoing, unsettled debate, we compare derived thresholds only within cohorts employing the same assay.

In the case of the two unsupervised learning methods (i.e., GMM and K-means clustering), we selected the thresholds that best separated the two target clusters (i.e., normal and abnormal) without considering the clinical diagnosis stage of the participants. Furthermore, both methods correct for participant age by adding age as an additional dimension to the clustering. In contrast, ROC analysis, tertile, and mean ±2 SD require the diagnostic stages of participants to define thresholds. In the case of ROC analysis, we calculated Youden’s index (*sensitivity* + *specificity* − 1) and selected the threshold that yielded the highest Youden’s index, thereby indicating the best separation of AD and CU participants (Ossenkoppele *et al*. 2018, Delmotte *et al*. 2021). The tertile and mean ±2 SD methods rely on the biomarker distributions of CU participants to define thresholds (Soldan *et al*. 2019, Mattsson-Carlgren *et al*. 2020). To some cohorts, the diagnosis-reliant methods could not be applied due to an insufficient number of participants in either of the stages. Finally, to gain robust threshold estimates, we calculated the 95% Confidence Intervals (CIs) for each CSF biomarker threshold based on 1,000 bootstrap samples.

### 2.4 Evaluating the concordance of ATN profiles across cohorts

To investigate how comparable participants of different cohorts were within each ATN profile and whether profilings based on different, study-specific thresholds would exhibit similar disease patterns, we performed hierarchical clustering on participants from all cohorts within each ATN biomarker profile. Low participant comparability would be indicated by a clustering that would be highly dominated by the dataset origin of participants, meaning participants of the same cohort cluster together, and thus cluster labels correlate significantly with cohort membership. Conversely, an evenly spread-out distribution of dataset members across clusters would indicate that similar disease patterns could be found across all cohorts’ participants in a respective ATN profile despite the differences in thresholds used for initial profile assignment.

Seven cohorts could be included in this analysis (ADNI, ARWIBO, DOD-ADNI, EDSD, JADNI, NACC, and PharmaCog), as finding a sufficient variable overlap between more cohorts proved infeasible. The clustering was based on 105 shared variables identified using ADataViewer (Salimi *et al*. 2022), 104 of which were MRI-measured brain region volumes with the remaining one being the Apolipoprotein E (APOE ε4) status **(Table S5)**. We deliberately omitted clinical assessments to mirror the biological perspective of the ATN framework. In order to adjust for the presence of potential batch effects, we applied the pyComBat correction on MRI variables (Behdenna *et al*. 2020). Cluster distances were calculated using Ward-linkage. The complete analysis was performed using K-Means and GMM respectively to gain the thresholds for profiling participants. ATN profiles with a total number of participants lower than 20 were excluded from this analysis (i.e., A-T+N-). To measure the association between dataset membership and assigned cluster labels, we calculated Cramer’s V with bias correction (Bergsma 2013) and assessed statistical significance using a chi^2^-test assuming a confidence level of 95%.

We repeated the hierarchical clustering using two procedures for determining the sought-after number of clusters per ATN profile. First, we optimized the number of clusters by calculating the cluster silhouette index across a range of possible clusterings and selected the number of clusters that maximized this index per ATN profile. In a second run, we assigned the number of clusters equal to the number of cohorts considered for each profile.

To provide an additional qualitative perspective, we applied the UMAP algorithm (McInnes *et al*. 2018) to all participants of the previously mentioned seven cohorts who were assigned to each ATN profile. UMAP projects a dataset into a lower dimensional space (here, two dimensions), while trying to preserve the global structure of the data. For this, we used the same 105 variables as in the clustering (**Table S5**). Again, ATN profiles with less than 20 participants (i.e., A-T+N-) were removed from this analysis. The resulting low-dimensional visualization provides a notion of whether participants assigned to the same ATN profile could be easily separated by cohort membership.

### 2.5 Data availability

All datasets used in this work are publicly accessible after successful access application. Links guiding to each individual resource can be found at: https://adata.scai.fraunhofer.de/cohorts.

## 3. Results

In this section, we first describe identified differences in CSF thresholds obtained by applying five commonly used thresholding approaches on 11 distinct cohort datasets. In this context, we further investigate how statistically conservative methods are and assess the uncertainty of their threshold estimates. Secondly, we compare the data-driven thresholds across subsets of cohorts that used the same CSF assay. Thirdly, we highlight how the difference in threshold values impacts the assignment of individuals to distinct ATN profiles. Finally, we explored the comparability and generalizability of biological patterns observed in ATN profiles across cohorts when their respectively different data-derived thresholds were applied.

### 3.1 CSF thresholds are method dependent

#### 3.1.1 Robustness of data-driven thresholds achieved using different methods

**Table 2** shows the CSF biomarker thresholds obtained for each cohort and thresholding method. When comparing the values yielded by applying different thresholding approaches to each cohort, we observed considerable differences among them. As an illustration, the thresholds obtained for Aβ1-42 in the ADNI cohort ranged from 280.8 (mean ±2 SD) to 983.7 (GMM). Another extreme case was observed in the EMIF_ELISA subgroup (i.e., the subgroup of EMIF participants with CSF values measured using ELISA), where the thresholds for tTau varied from 269.4 (tertile) to 669.4 (mean ±2 SD). In general, the largest difference across estimated threshold values was found for Aβ1-42 with an average deviation of 66%, calculated with respect to each cohort’s largest observed Aβ1-42 threshold **(Table S8)**. For tTau and pTau, we discovered average deviations of 46% and 49%, respectively.

**Table 2:** Thresholds obtained for each CSF biomarker using different data-driven methods. ‘**-’:** Method application infeasible due to data limitations. **Note:** Cohorts are grouped together based on the used CSF assay and are separated by thick lines. The NACC and EMIF cohorts were divided into separate groups based on the CSF assay used for measuring the CSF biomarkers within each cohort. All values are given as pg/mL. In order for Aβ1-42 measurements to be considered abnormal, the measurement needs to place below the threshold, while for Tau biomarkers an abnormal value exceeds the threshold.

| Cohort | Method |  |  |  |  |  |  |  |  |  |  |  |  |  |  |
| --- | --- | --- | --- | --- | --- | --- | --- | --- | --- | --- | --- | --- | --- | --- | --- |
| | GMM | | | K-means | | | Tertile | | | ROC | | | Mean ±2 SD | | |
| | Aβ1-42 | pTau | tTau | Aβ1-42 | pTau | tTau | Aβ1-42 | pTau | tTau | Aβ1-42 | pTau | tTau | Aβ1-42 | pTau | tTau |
| ADNI | 983.7 | 34.3 | 352.3 | 973 | 28.1 | 286.2 | 832.1 | 21.5 | 240.5 | 801.7 | 24.2 | 264.8 | 280.8 | 40.9 | 409.7 |
| EPAD | 1034.9 | 27.7 | 306.7 | 1030.7 | 19.8 | 223.7 | 883 | 19.9 | 228.3 | 731.5 | 20.9 | 213.8 | 337.9 | 43.3 | 436.4 |
| AIBL | 714.4 | 92.6 | 591.5 | 689.9 | 77.7 | 635.3 | - | - | - | - | - | - | - | - | - |
| ARWIBO | 577.5 | 126.2 | 499 | 512.8 | 69.2 | 406.2 | - | - | - | - | - | - | - | - | - |
| EDSD | 737.5 | 113.2 | 535.7 | 790.8 | 82.8 | 577.7 | - | - | - | - | - | - | - | - | - |
| PharmaCog | 779.6 | 93.1 | 630.6 | 773.6 | 66.2 | 465.8 | - | - | - | - | - | - | - | - | - |
| PREVENT-AD | 1030.9 | 68 | 424.8 | 1150.4 | 50.3 | 301.6 | 1104.4 | 53.1 | 304.2 | - | - | - | 588.4 | 89.6 | 598.1 |
| NACC_ELISA | 596.2 | 80.8 | 537.4 | 591.4 | 61.1 | 497.4 | 629 | 48 | 376.6 | 474 | 64 | 468 | 291.5 | 88 | 728.8 |
| EMIF_ELISA | 730.9 | 81.4 | 523.8 | 723.7 | 65 | 435.1 | 538.2 | 52.1 | 269.4 | 560 | 59 | 355.4 | 173.1 | 95.1 | 669.4 |
| NACC_xMAP | 298.6 | 58.4 | 94 | 270.3 | 40.2 | 70 | 244.8 | 36.3 | 53.8 | 226.4 | 41.2 | 56.7 | 21.8 | 69.3 | 99.7 |
| EMIF_xMAP | 400.4 | 51.7 | 204.2 | 356.4 | 37.6 | 129.9 | 450.2 | 30.8 | 83.2 | 352.2 | 28.7 | 100.1 | 239.5 | 55.2 | 123.4 |
| DOD-ADNI | 1276 | 26.5 | 306.1 | 1231.9 | 22.5 | 250.2 | 927 | 19.8 | 229 | - | - | - | 282.4 | 34.3 | 369.9 |
| JADNI | 416.4 | 61.3 | 133.8 | 414.3 | 62 | 145.8 | 392 | 38.9 | 72.2 | 333.9 | 44.9 | 86.2 | 186.9 | 66.2 | 128.2 |

To further estimate the robustness for the threshold estimates, we repeated the thresholding using random subsets of a respective cohort’s participants and calculated 95 % CIs for the estimate **(Table S6)**. Across all cohorts and thresholding methods, the uncertainty of estimated values remained rather small (below 1% relative difference with respect to the upper CI bound) with a maximum relative difference of 8.9% for the pTau threshold of ARWIBO estimated using GMM (95% CI [148-162]; **Table S7**). Surprisingly, only 42 of the 153 thresholds obtained on the full cohort datasets placed within their corresponding estimated CI, however, we found that most deviations from the respectively upper or lower bound were numerically small and clinically neglectable (e.g., ADNI tTau threshold estimated using K-means of 286.2 versus a 95% CI of [288 - 289]). Exceptionally large deviations could still be found, such as, for example, the GMM-derived threshold for Aβ1-42 in PREVENT_AD (full cohort point estimate: 1031; 95% CI: [1159-1216], or the ROC-determined Aβ1-42 value in EPAD (full cohort: 732; 95% CI: [681-689]).

In general, we found that the most conservative profiling thresholds (i.e., yielding numerically more extreme thresholds that make it less likely for biomarker measurements to be considered abnormal) were obtained using the mean ±2 SD method **(Table 2; Figure 1)**, while the clustering-based approaches (K-means and GMM) and ROC analysis often estimated thresholds that were less conservative, and the tertile method usually was the least conservative.

**Figure 1:**
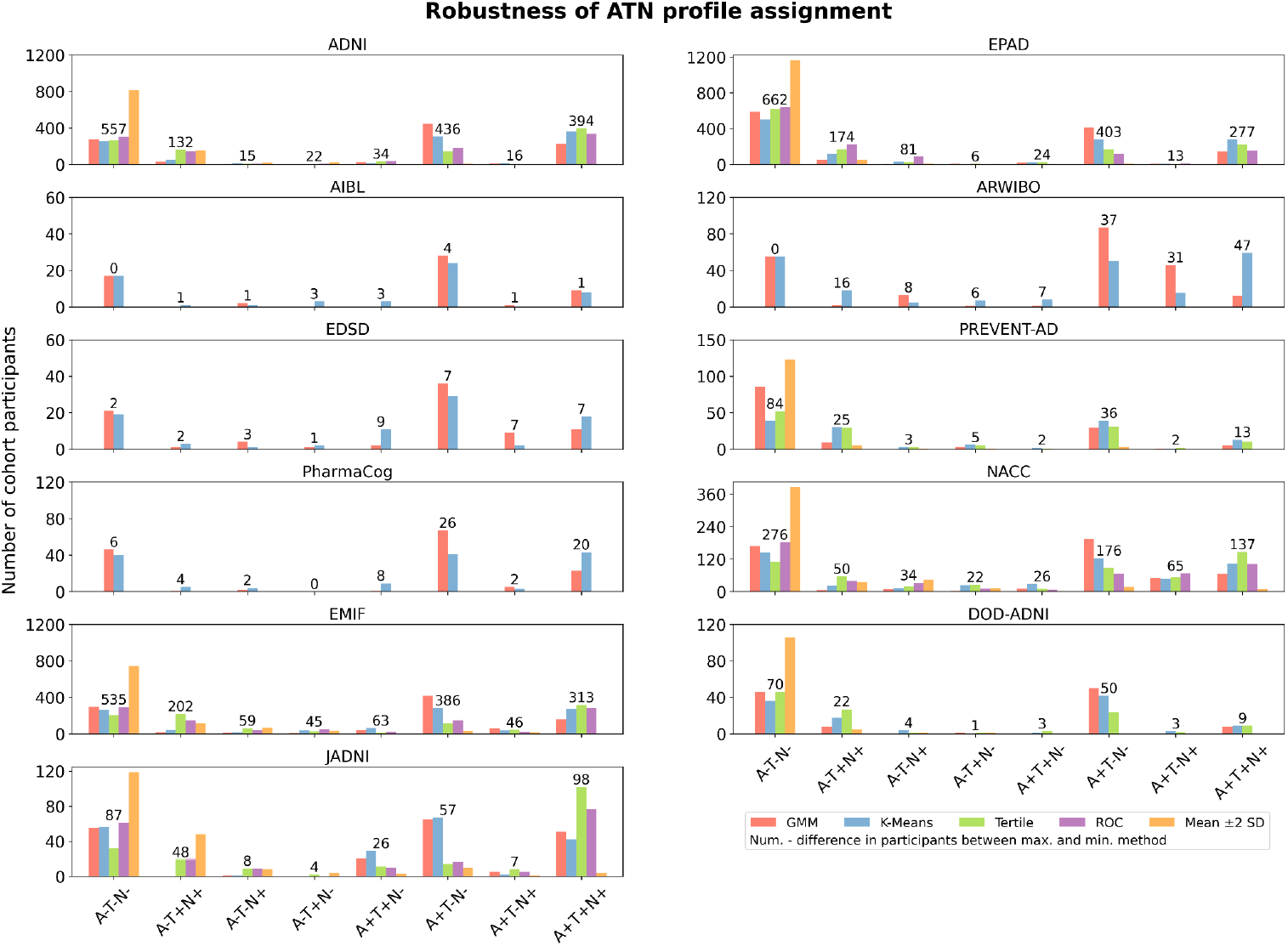
Proportion of participants categorized to each ATN profile by the different thresholding techniques across cohorts. The y-axis shows the proportion of participants assigned to each of the eight biomarker profiles in each cohort. The absolute number of patients assigned for each category is displayed in Supplementary Tables S11-15.

#### 3.1.2 Evaluating the robustness of data-driven thresholds across cohorts

As some studies used common assays, we investigated whether the thresholds we obtained using data-driven methods would be similar across cohorts that employed the same assay. The majority of thresholds varied substantially from one another, although some similarities were observed (**Table 2**). For instance, the thresholds obtained for cohorts using the Roche Elecsys immunoassay (ADNI and EPAD) were quite similar across biomarkers with an average difference of 8%, 15%, and 13% of the corresponding highest value of Aβ1-42, pTau, and tTau, respectively **(Table S9)**. Among the seven cohorts using the INNOTEST® assays, the average across-cohort difference of Aβ1-42, pTau, and tTau thresholds respectively amounted to 47%, 22%, and 31% of the highest value per biomarker. Here, especially the Aβ1-42 values of PREVENT-AD showed substantial deviations from values gained on other cohorts **(Table 2)**. Lastly, among the four cohorts employing a multiplex xMAP assay, we found average differences of 71%, 51%, and 67% for Aβ1-42, pTau, and tTau, respectively.

When investigating which of the five investigated thresholding methods obtained the most consistent estimates (i.e., minimal deviation) across cohorts and biomarkers, we found that the tertile method was most consistent for the Roche Elecsys immunoassay datasets (average difference across cohorts and biomarkers of 6%) **(Table S10)**. Further, thresholds estimated using the ROC analysis showed the lowest deviation for the INNOTEST® assay-employing cohorts, as well as for the cohorts using a multiplex xMAP assay with a respective average difference of 16% and 38% across cohorts and biomarkers.

### 3.2 Impact of thresholding method selection on ATN profiling

Next, we evaluated the impact that different thresholds defined using distinct thresholding methods had on the resulting ATN profiling. For this, we applied each method’s threshold and assessed the robustness of resulting participant assignments (Numbers of samples per cohort, method, and ATN profile are provided in **Tables S11-S15**). Unsurprisingly, given the magnitude of observed differences **(Table 2)**, method selection substantially influenced the achieved profiling of participants, as vast differences in the number of participants per ATN profile were identified across applied methods **(Figure 1)**. In alignment with the previously presented results, depending on how conservative the thresholding methods were, fewer or more participants were assigned to ATN profiles that involved abnormal biomarker measurements. Here, the largest gaps in participant assignments between methods involved the mean ±2 SD approach, which assigned most participants to the A-T-N-profile. In contrast, the two clustering techniques often returned quite similar participant counts, while the tertile method assigned relatively more individuals to the A+T+N+ profile. Large gaps between the distinctly thresholded participant assignments were, for example, observed in ADNI, where differences between the highest and lowest participant counts of profiles amounted to 557 individuals for A-T-N-, 132 for A-T+N+, 436 for A+T-N-, and 394 for A+T+N+ with respect to a total of 1017 ADNI participants. Nonetheless, variations strongly depended on the underlying cohort and in some biomarker profiles and cohorts an interpretation of the observed numbers was difficult due to the limited sample size (e.g., AIBL).

### 3.3 Generalizability of ATN profile-specific biological signals across cohorts

We evaluated whether individuals of disparate cohorts assigned to the same ATN profile exhibited comparable disease patterns and, thereby, if ATN-based results could be generalized. To this end, we conducted a hierarchical clustering of participants from seven cohorts within each ATN profile based on 105 shared variables (104 MRI measurements and APOE ε4 allele status). The ATN profiles were defined using both K-means and GMM, respectively. Afterward, the correlation between assigned cluster labels and cohort membership was assessed to evaluate if individuals cluster together based on biological signals or shared cohort origin.

When determining the optimal number of clusters for partitioning participants, we found that only two or three clusters already provided the best clustering solution (**Figure S2**). Sample sizes for each cluster and profile are provided in **Tables S18-S31**. We identified no significant associations between cluster labels and cohort membership in ATN profiles defined using K-means (p>0.05; **Table 3**), indicating that the clustering was not dominated by individuals’ cohort origin. Only when applying GMM for thresholding, a significant and strong association was discovered in the A+T+N-profile (Cramer’s V=0.48, p<0.01), as well as a weaker association in the A+T+N+ profile (Cramer’s V=0.26, p=0.04).

**Table 3:**
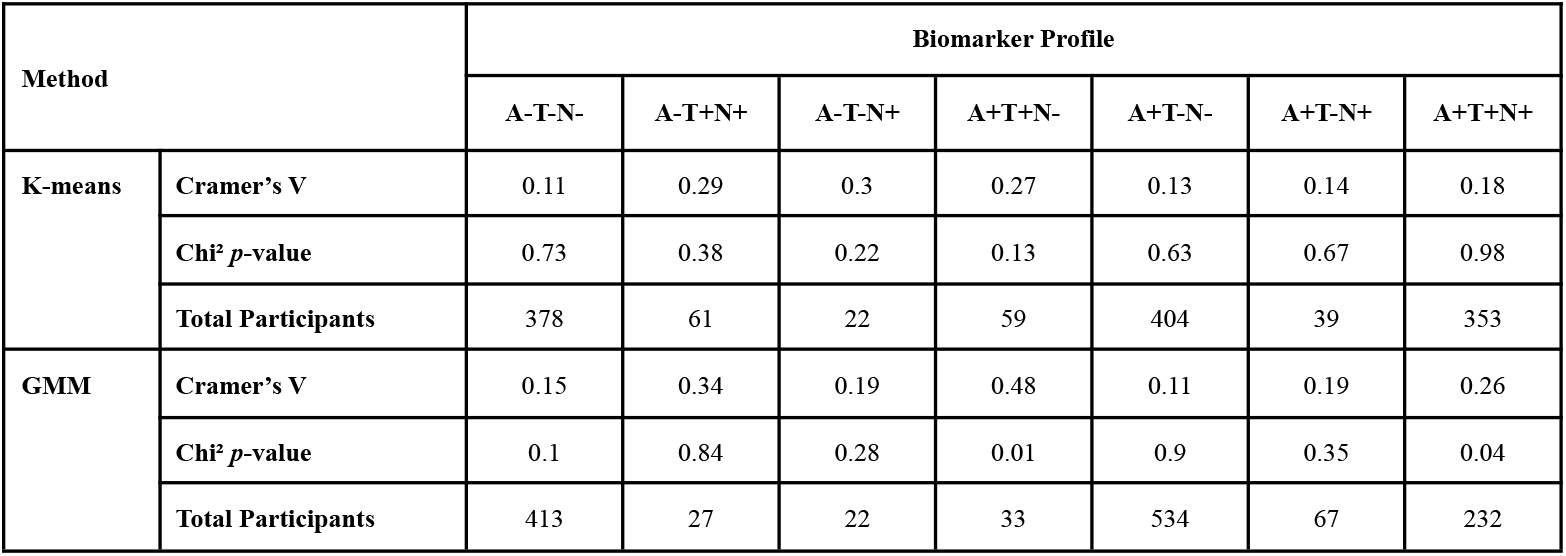
The Cramer’s V and the p-value for each clustering of participants in each ATN profile using certain data-driven thresholds. **Note:** The ATN profile with less than 20 participants was excluded (i.e., A-T+N-).

In a second variation of the analysis, we set the number of clusters equal to the number of cohorts per ATN profile (contingency tables are presented in **Tables S33-S46**). We did not observe any significant associations for both K-means and GMM-defined ATN profiles, respectively **(Table S32)**.

To provide a visual intuition about how similar individuals of the same ATN profile were across cohorts, we generated a UMAP visualization based on the same cohorts and variables used in the previously mentioned clustering approach. The total number of participants included in this analysis per cohort are presented in **Tables S16-S17**. While interpreting the absolute distances in a UMAP visualization is futile, the grouping of individuals from different cohorts and the relative distances between participants of the same cohort dataset compared to the distances found between participants from independent cohorts provide an indication of their respective similarity.

In ATN profiles defined using both K-means **(Figure 2A)** and GMM **(Figure 2B)** respectively, we observed individuals from multiple cohorts in all visible groupings. Simultaneously, members of each cohort distributed widely across the UMAP space and were often positioned closer to other datasets’ participants than to their peers. Especially in the more populated biomarker profiles (A-T-N-, A+T-N-, and A+T+N+), we observed no clear separation of participants by dataset origin. A thorough interpretation of sparser profiles, however, remained difficult due to the unequal and sometimes low sample sizes of cohorts. Conclusively, both the clustering approach and UMAP projection were largely not governed by cohort membership, but by biological signals as expressed predominantly through MRI variables.

**Figure 2:**
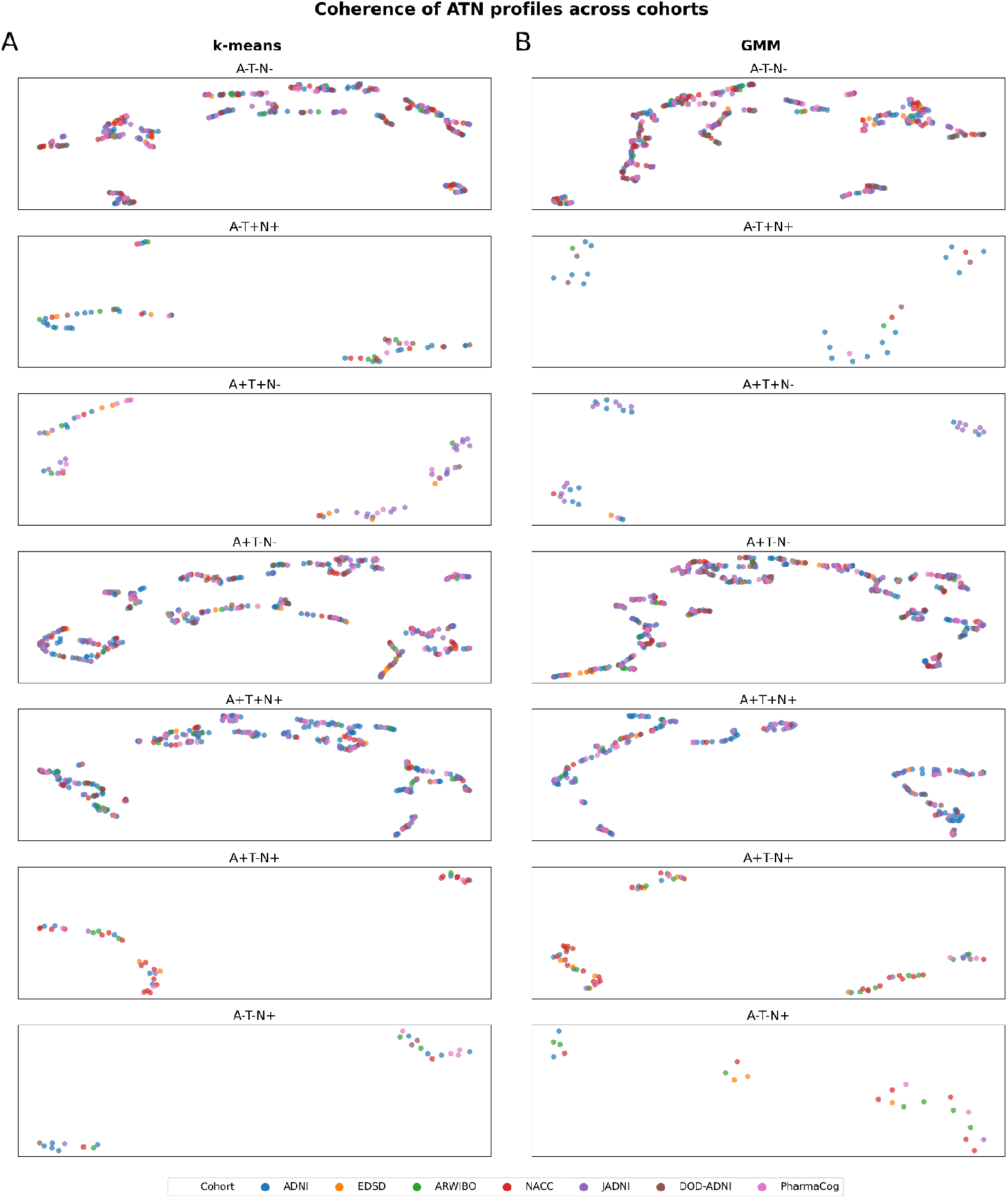
Identification of potential participants’ subgroups categorized in each ATN profile among cohorts using UMAP. **A)** ATN profiles achieved using K-means thresholds. **B)** ATN profiles achieved using GMM thresholds. **Note:** Missing profiles were removed due to the lack of participants (number of participants below 20). No axis labeling is provided as they are not directly interpretable. The total number of participants in each ATN profile are presented in supplementary **Tables S16-S17**.

## 4. Discussion

In this work, we investigated the robustness and generalizability of the ATN framework across eleven AD cohort datasets and five commonly used data-driven approaches for defining biomarker thresholds. When comparing thresholds yielded by distinct methods for the same biomarker, we observed substantial variation that showcased the contrasting statistical properties of thresholding methods and the impact that method selection has on achieved ATN profilings. Even when CSF biomarkers were measured using the same assay, applying the same thresholding method led to estimates that often deviated substantially across cohorts. These findings indicated that thresholds are most often not interchangeable across cohorts without disrupting the ATN profile assignments of individuals. Further, it is not certain that independent data-driven profilings achieved on disparate cohort datasets always capture comparable individuals exhibiting similar disease patterns in the same ATN profile. However, clustering participants of seven cohorts within each ATN profile revealed no clear cluster separation by dataset membership, indicating that participants with similar MRI-based biological signals could be identified from individually profiled cohorts. This showed that patterns discovered in independently ATN profiled cohorts could still be comparable even though numerical differences existed in the respectively applied thresholds.

### 4.1 Determined thresholds are dataset and method specific

Determining thresholds for Aβ1-42, pTau, and tTau by applying five established thresholding methods across AD datasets revealed large variations depending on which method and dataset were used. We speculate that these differences likely find their origin in the initial exclusion and exclusion criteria of cohort studies that define the population from which their participants were recruited (Salimi *et al*. 2022), as well as variation in biomarker assessment (Hampel *et al*. 2021, Dumurgier *et al*. 2013). Additionally, the identified differences could be promoted by assumptions that thresholding methods make on the data from which values are estimated. For example, when applying methods that rely on the clinical diagnosis of participants, thresholds could be distinctly influenced by discrepant diagnostic criteria or differences in the average disease stage of participants between two cohorts. The AD diagnosis of PharmaCog participants, for example, was defined based on amnestic MCI with low Aβ1-42 CSF levels (Galluzzi *et al*. 2016), while participants in AIBL were diagnosed using the NINCDS-ADRDA criteria (Ellis *et al*. 2009).

Whether thresholds are interchangeable between cohorts depends strongly on the data they were estimated from as well as the applied thresholding method. While previous studies have found that thresholds were interchangeable for their investigated pair of cohorts (Hansson *et al*. 2018, Ingala *et al*. 2021), we observed no general, overarching consensus across eleven cohorts even in subgroups where the same assay was used to measure biomarker values. Applying a threshold derived from one cohort to another cohort will flip the +/- labels of participants that are in between the transferred threshold and the other cohort’s original threshold. Effectively, this means that with increasing distance between thresholds, more individuals will be assigned to other profiles if these thresholds were transferred. Therefore, thresholds could be interchanged with a lower impact on the resulting ATN profiling between ADNI and EPAD, yet, applying PharmaCog’s threshold of tTau on PREVENT-AD would cause considerable disruption of the original PREVENT-AD profiles. This also raises doubts about the common practice to take thresholds estimated on different datasets to profile participants in a new cohort at hand (Ebenau et al. 2020).

### 4.2 Statistical implications of thresholding method selection

The assumptions made about the data-underlying distributions differ vastly across the five evaluated thresholding methods and form their respective statistical properties. Employing a thresholding method that is more conservative (e.g., mean ±2 SD) will produce an ATN profiling of individuals where fewer individuals will be assigned to profiles with abnormal biomarker measurements, while less conservative methods (e.g., tertile or GMM) will do the opposite. This directly links the method selection to the probability of committing type I (i.e., biomarker falsely considered abnormal) and type II (i.e., biomarker falsely considered normal) errors in our resulting profiling and implies that it will inevitably introduce some form of statistical bias into analyses performed on the achieved ATN profiles. From our experiments, we observed that such statistical properties are not only of theoretical interest but hugely impact the assignment of participants, as it solely depends on the determined threshold. Therefore, a meaningful comparison of ATN profiles and, consequently, achieved results across studies is likely futile if they relied on different thresholding methods.

Furthermore, in a statistical sense, thresholds themselves are point estimates that are made with uncertainty. We used bootstrapping to assess the uncertainty of our calculated thresholds which revealed relatively stable estimates across most methods and biomarkers. However, in rarer cases, the threshold estimates made based on the complete dataset varied substantially from our bootstrapped CIs which raises concerns about the profilings that would result from them. Consequently, careful uncertainty estimation for data-driven thresholds is imperative before employing them in analyses, yet, such uncertainty estimation was seldom done in studies utilizing the ATN framework. Only 11 of the publications we reviewed for commonly applied thresholding methods mentioned some form of uncertainty estimation.

### 4.3 Results achieved on ATN profiles derived from single cohorts might generalize

A yet underexplored aspect of the ATN framework was the generalizability of results gained on cohort-specific profilings. As thresholds are data-derived, it is possible that not only their numerical values differ but also that comparability of participants assigned to the same ATN profile is not guaranteed across cohorts.

Using a clustering approach, we observed that participants of all cohorts formed mixed clusters within each respective ATN profile. Only in two profiles (A+T+N- and A+T+N+), a significant association between cohort membership and cluster assignment was identified, when thresholds were defined using GMM. We believe our results show that ATN-based findings can generalize across different cohorts’ ATN profilings in principle, yet utmost care should be paid to this aspect. As our results are based on a clustering approach and predominantly MRI data, it remains very well possible that the generalizability of results could be data modality dependent and might vary for different analyses types such as data-driven prognosis or diagnosis. However, one considerable limitation in generalizability can be caused by differing thresholding methods. Discoveries made in a study that employed a very conservative threshold established, for example, using mean ±2 SD are unlikely to be validated in a study relying on GMM for thresholding. Conclusively, in an ATN-based setting, it is especially important that results are externally validated in independent data sources while keeping threshold characteristics in mind.

### 4.5 Limitations

One limitation of our study is that only CSF biomarkers were used and that further suggested biomarkers such as PET imaging (Jack *et al*. 2018) were not investigated due to their limited availability across datasets. Previous work found that fluid and imaging biomarkers were not always concordant and appeared to be stage-dependent (Mattsson-Carlgren *et al*. 2020, Illán-Gala *et al*. 2018), thus, our results might be CSF biomarker specific. Further limitations can be seen in the comparably low sample sizes in some ATN profiles, which partially result due to the natural progression of AD as well as already limited numbers of participants providing CSF samples in the complete cohorts. Finally, missing clinical diagnoses prohibited the application of some thresholding methods in a few cohorts.

### 4.6 Conclusion

The introduction of the ATN framework constitutes an important step toward a biologically profound definition of AD that differentiates it from other dementias. However, it is crucial that the intricacies and pitfalls of ATN profiling are well understood by research practitioners in order to generate reliable and robust insights that generalize beyond their discovery data. The results presented in this work highlight that the selection of any specific data-driven thresholding method represents a decision that will inevitably introduce a statistical bias into obtained ATN profiles which will also propagate to subsequent analyses. A specific ATN profile thresholded using a highly conservative method is not equal to one defined by, for example, a less conservative clustering approach. Thereby, they should not be regarded as equal although their assigned label (i.e., A+T+N-) implies this. Further, while interchanging biomarker thresholds across cohorts was feasible in single examples (Hansson *et al*. 2018), from a broader perspective across many cohorts it seems improbable that this holds for any two datasets and thresholding methods. Finally, when comparing clusters based on MRI signals exhibited in each respective ATN profile across cohorts, we only identified significant differences among distinct datasets in two profiles of one thresholding method. This could imply that numerical differences in thresholding values might play a lesser role as long as they are applied to the data they were estimated on and cohorts share similar recruitment criteria. We want to emphasize, however, that this constitutes no ‘carte blanche’ for considering all signals equal across individuals assigned to the same ATN profile based on different data. The properties of the data, cohorts’ selection criteria, and employed thresholding method need to be thoroughly investigated whenever data-driven thresholding approaches are used. Establishing harmonized, validated, and potentially experimentally-derived thresholds for each ATN biomarker would circumvent many of the pitfalls in data-driven estimation and substantially improve the generalizability of ATN-based results (Hampel *et al*. 2021).

## Abbreviations

Aβ1-42: Amyloid-β 1-42
AD: Alzheimer’s disease
ADNI: Alzheimer’s Disease Neuroimaging Initiative
AIBL: Australian Imaging, Biomarker & Lifestyle Flagship Study of Ageing
ARWIBO: Alzheimer’s Disease Repository Without Borders
ATN: Amyloid, Tau, and Neurodegeneration
PET: Positron emission tomography
CDR: Clinical Dementia Rating
CIs: Confidence Intervals
CSF: Cerebrospinal fluid
CU: Cognitively unimpared
DOD-ADNI: Effects of TBI & PTSD on Alzheimer’s Disease in Vietnam Vets
EDSD: European DTI Study on Dementia
EMIF: European Medical Information Framework
EPAD: European Prevention of Alzheimer’s Dementia
GMMs: Gaussian mixture models
JADNI: Japanese Alzheimer’s Disease Neuroimaging Initiative
MCI: Mild cognitive impairment
MMSE: Mini-Mental State Examination
MRI: Magnetic resonance imaging
NACC: National Alzheimer’s Coordinating Center
NINCDS-ADRDA: National Institute of Neurological and Communicative Diseases and Stroke/Alzheimer’s Disease and Related Disorders Association
PharmaCog: Prediction of Cognitive Properties of New Drug Candidates for Neurodegenerative Diseases in Early Clinical Development
PREVENT-AD: Pre-symptomatic Evaluation of Experimental or Novel Treatments for Alzheimer’s Disease
pTau: Phosphorylated tau
ROC: Receiver operating characteristic
SD: Standard deviation
tTau: Total tau

## Acknowledgments

We want to commend all data owners on their adherence to open science principles by sharing their data. We believe that their commitment is invaluable for any scientific research.

Data collection and sharing of ARWIBO was supported by the Italian Ministry of Health, under the following grant agreements: Ricerca Corrente IRCCS Fatebenefratelli, Linea di Ricerca 2; Progetto Finalizzato Strategico 2000-2001 “Archivio normativo italiano di morfometria cerebrale con risonanza magnetica (età 40+)”; Progetto Finalizzato Strategico 2000-2001 “Decadimento cognitivo lieve non dementigeno: stadio preclinico di malattia di Alzheimer e demenza vascolare. Caratterizzazione clinica, strumentale, genetica e neurobiologica e sviluppo di criteri diagnostici utilizzabili nella realtà nazionale,”; Progetto Finalizzata 2002 “Sviluppo di indicatori di danno cerebrovascolare clinicamente significativo alla risonanza magnetica strutturale”; Progetto Fondazione CARIPLO 2005-2007 “Geni di suscettibilità per gli endofenotipi associati a malattie psichiatriche e dementigene”; “Fitness and Solidarietà”; and anonymous donors.

EPAD LCS is registered at www.clinicaltrials.gov Identifier: NCT02804789. Data used in preparation of this article were obtained from the EPAD LCS data set V.IMI, doi:10.34688/epadlcs_v.imi_20.10.30. The EPAD LCS was launched in 2015 as a public private partnership, led by Chief Investigator Professor Craig Ritchie MB BS. The primary research goal of the EPAD LCS is to provide a well-phenotyped probability-spectrum population for developing and continuously improving disease models for Alzheimer’s disease in individuals without dementia. This work used data and/or samples from the EPAD project which received support from the EU/EFPIA Innovative Medicines Initiative Joint Undertaking EPAD grant agreement n° 115736 and an Alzheimer’s Association Grant (SG21-818099-EPAD).

PharmaCog was funded through the European Community’s ‘Seventh Framework’ Programme (FP7/2007-2013) for an innovative scheme, the Innovative Medicines Initiative (IMI). IMI is a young and unique public-private partnership, founded in 2008 by the pharmaceutical industry (represented by the European Federation of Pharmaceutical Industries and Associations), EFPIA and the European Communities (represented by the European Commission).

J-ADNI was supported by the following grants: Translational Research Promotion Project from the New Energy and Industrial Technology Development Organization of Japan; Research on Dementia, Health Labor Sciences Research Grant; Life Science Database Integration Project of Japan Science and Technology Agency; Research Association of Biotechnology (contributed by Astellas Pharma Inc., Bristol-Myers Squibb, Daiichi-Sankyo, Eisai, Eli Lilly and Company, Merck-Banyu, Mitsubishi Tanabe Pharma, Pfizer Inc., Shionogi & Co., Ltd., Sumitomo Dainippon, and Takeda Pharmaceutical Company), Japan, and a grant from an anonymous Foundation.

Data collection and sharing for this project was funded by the Alzheimer’s Disease Neuroimaging Initiative (ADNI) (National Institutes of Health Grant U01 AG024904) and DOD ADNI (Department of Defense award number W81XWH-12-2-0012). ADNI is funded by the National Institute on Aging, the National Institute of Biomedical Imaging and Bioengineering, and through generous contributions from the following: AbbVie, Alzheimer’s Association; Alzheimer’s Drug Discovery Foundation; Araclon Biotech; BioClinica, Inc.; Biogen; Bristol-Myers Squibb Company; CereSpir, Inc.; Cogstate; Eisai Inc.; Elan Pharmaceuticals, Inc.; Eli Lilly and Company; EuroImmun; F. Hoffmann-La Roche Ltd and its affiliated company Genentech, Inc.; Fujirebio; GE Healthcare; IXICO Ltd.; Janssen Alzheimer Immunotherapy Research & Development, LLC.; Johnson & Johnson Pharmaceutical Research & Development LLC.; Lumosity; Lundbeck; Merck & Co., Inc.; Meso Scale Diagnostics, LLC.; NeuroRx Research; Neurotrack Technologies; Novartis Pharmaceuticals Corporation; Pfizer Inc.; Piramal Imaging; Servier; Takeda Pharmaceutical Company; and Transition Therapeutics. The Canadian Institutes of Health Research is providing funds to support ADNI clinical sites in Canada. Private sector contributions are facilitated by the Foundation for the National Institutes of Health (www.fnih.org). The grantee organization is the Northern California Institute for Research and Education, and the study is coordinated by the Alzheimer’s Therapeutic Research Institute at the University of Southern California. ADNI data are disseminated by the Laboratory for Neuro Imaging at the University of Southern California. This research was also supported by NIH grants P30 AG010129 and K01 AG030514.

The NACC database is funded by NIA/NIH Grant U01 AG016976. NACC data are contributed by the NIA-funded ADCs: P30 AG019610 (PI Eric Reiman, MD), P30 AG013846 (PI Neil Kowall, MD), P30 AG062428-01 (PI James Leverenz, MD) P50 AG008702 (PI Scott Small, MD), P50 AG025688 (PI Allan Levey, MD, PhD), P50 AG047266 (PI Todd Golde, MD, PhD), P30 AG010133 (PI Andrew Saykin, PsyD), P50 AG005146 (PI Marilyn Albert, PhD), P30 AG062421-01 (PI Bradley Hyman, MD, PhD), P30 AG062422-01 (PI Ronald Petersen, MD, PhD), P50 AG005138 (PI Mary Sano, PhD), P30 AG008051 (PI Thomas Wisniewski, MD), P30 AG013854 (PI Robert Vassar, PhD), P30 AG008017 (PI Jeffrey Kaye, MD), P30 AG010161 (PI David Bennett, MD), P50 AG047366 (PI Victor Henderson, MD, MS), P30 AG010129 (PI Charles DeCarli, MD), P50 AG016573 (PI Frank LaFerla, PhD), P30 AG062429-01(PI James Brewer, MD, PhD), P50 AG023501 (PI Bruce Miller, MD), P30 AG035982 (PI Russell Swerdlow, MD), P30 AG028383 (PI Linda Van Eldik, PhD), P30 AG053760 (PI Henry Paulson, MD, PhD), P30 AG010124 (PI John Trojanowski, MD, PhD), P50 AG005133 (PI Oscar Lopez, MD), P50 AG005142 (PI Helena Chui, MD), P30 AG012300 (PI Roger Rosenberg, MD), P30 AG049638 (PI Suzanne Craft, PhD), P50 AG005136 (PI Thomas Grabowski, MD), P30 AG062715-01 (PI Sanjay Asthana, MD, FRCP), P50 AG005681 (PI John Morris, MD), P50 AG047270 (PI Stephen Strittmatter, MD, PhD).

## Author contributions

CB conceived the study. CB and YS collected the datasets. YS prepared the datasets and conducted the analysis. DDF contributed to the analysis. CB, YS, and DDF wrote the manuscript. MHA revised the manuscript and acquired the funding.

