## Supplementary Material for "Exploring the intricacies and pitfalls of the ATN framework: An assessment across cohorts and thresholding methodologies"

| Cohort | Aβ1-42 | pTau | tTau |
| --- | --- | --- | --- |
| **ADNI** | <200, >1700 | <8, >120 | <80, >1300 |
| **ARWIBO** | - | <15.6 | <75, >1200, >1209.5 |
| **EPAD** | <200, >1700 | <8 | <80 |

**Table S1:** Technical limit for the CSF biomarkers reported in certain cohort studies. We excluded the participants with measured CSF below or above the technical limit values.

| Cohort | # Patients | Age | Female % | Education | APOE 4 % | CDR | MMSE | CDRSB | Hippocampus | tTau | pTau | Aβ1-42 |
| --- | --- | --- | --- | --- | --- | --- | --- | --- | --- | --- | --- | --- |
| **ADNI [1]** | 1017 | 73.5 (7.3) | 43.5 | 16.0 (2.7) | 52.3 | 0.4 (0.3) | 27.1 (2.7) | 1.8 (1.8) | 6756.7 (1173.0) | 290.0 (135.8) | 28.4 (15.1) | 848.3 (364.5) |
| **EPAD [2]** | 1237 | 66.1 (7.4) | 53.9 | 14.4 (3.7) | 47.4 | 0.2 (0.2) | 28.3 (2.1) | 0.4 (0.8) | 4730.3 (750.8) | 225.6 (107.3) | 20.1 (11.8) | 1056.4 (362.9) |
| **AIBL [3]** | 57 | 73.8 (6.4) | 45.6 | 12.7 (2.8) | 34.5 | 0.3 (0.4) | 26.4 (4.9) | 1.6 (2.7) | 2.8 (0.4) | 438.8 (276.1) | 68.5 (30.5) | 633.0 (241.8) |
| **ARWIBO [4]** | 217 | 70.5 (8.0) | 54.4 | 7.6 (4.0) | 39.5 | 0.7 (0.5) | 23.3 (4.8) | - | 6155.9 (1225.4) | 428.3 (255.7) | 71.9 (46.7) | 513.5 (239.2) |
| **EDSD [5]** | 86 | 70.0 (6.9) | 44.2 | 11.8 (3.0) | 54.7 | - | 26.6 (2.1) | - | 6995.7 (1178.0) | 456.8 (276.8) | 82.0 (40.6) | 665.1 (345.9) |
| **PREVENT-AD [6]** | 133 | 62.8 (5.3) | 67.7 | 15.2 (3.0) | - | - | - | - | - | 284.8 (156.6) | 49.0 (20.3) | 1184.9 (298.3) |
| **PharmaCog [7]** | 145 | 69.2 (7.3) | 57.2 | 10.6 (4.4) | 46.2 | 0.5 (0.0) | 26.6 (1.8) | - | 6729.3 (1420.6) | 475.5 (345.2) | 67.6 (34.7) | 693.0 (292.5) |
| **NACC [8]** | 506 | 67.4 (10.0) | 50.2 | 15.7 (3.1) | 46.6 | 0.4 (0.5) | 26.1 (4.8) | 2.0 (2.6) | 6.4 (0.9) | 235.0 (288.2) | 48.4 (34.9) | 389.9 (254.2) |
| **EMIF [9]** | 1014 | 67.9 (8.6) | 44.8 | 11.2 (4.1) | - | 0.4 (0.3) | 26.1 (3.8) | - | 6956.4 (1212.5) | 377.5 (327.6) | 60.0 (33.6) | 586.4 (282.6) |
| **DOD-ADNI [10]** | 113 | 68.5 (4.2) | 0.9 | 15.2 (2.3) | 25.7 | 0.1 (0.2) | 28.4 (1.5) | 0.4 (0.7) | 7741.6 (970.7) | 219.6 (80.8) | 19.1 (8.3) | 1242.6 (490.9) |
| **JADNI [11]** | 197 | 71.3 (6.8) | 50.8 | 13.4 (2.8) | 47.7 | 0.4 (0.3) | 26.3 (3.0) | 1.8 (1.7) | 6221.8 (1229.4) | 114.1 (61.2) | 56.0 (24.3) | 354.1 (146.5) |

**Table S2:** Summary statistics of participants in each cohort study. Numerical measurements are reported as mean and standard deviation in parentheses. Categorical variables are presented based on the proportion of participants within a category. Note: APOE4 %, the proportion of participants with at least one APOE e4 status.

| Cohort | # Participants | Assay |
|---|---|---|
| NACC | 205 | INNOTEST® kit assay (Innogenetics, Ghent, Belgium), Enzyme-linked immunosorbent assay (ELISA) |
| | 301 | Multiplex xMAP Luminex platform (LuminexCorp., Austin, TX, USA) with Innogenetics (INNO-BIA AlzBio3, Ghent, Belgium) immunoassay |
| EMIF | 811 | INNOTEST® kit assay (Innogenetics, Ghent, Belgium), Enzyme-linked immunosorbent assay (ELISA) |
| | 203 | Multiplex xMAP Luminex platform (LuminexCorp., Austin, TX, USA) with Innogenetics (INNO-BIA AlzBio3, Ghent, Belgium) immunoassay |

**Table S3:** Number of participants in certain cohorts with CSF measurements using each assay.

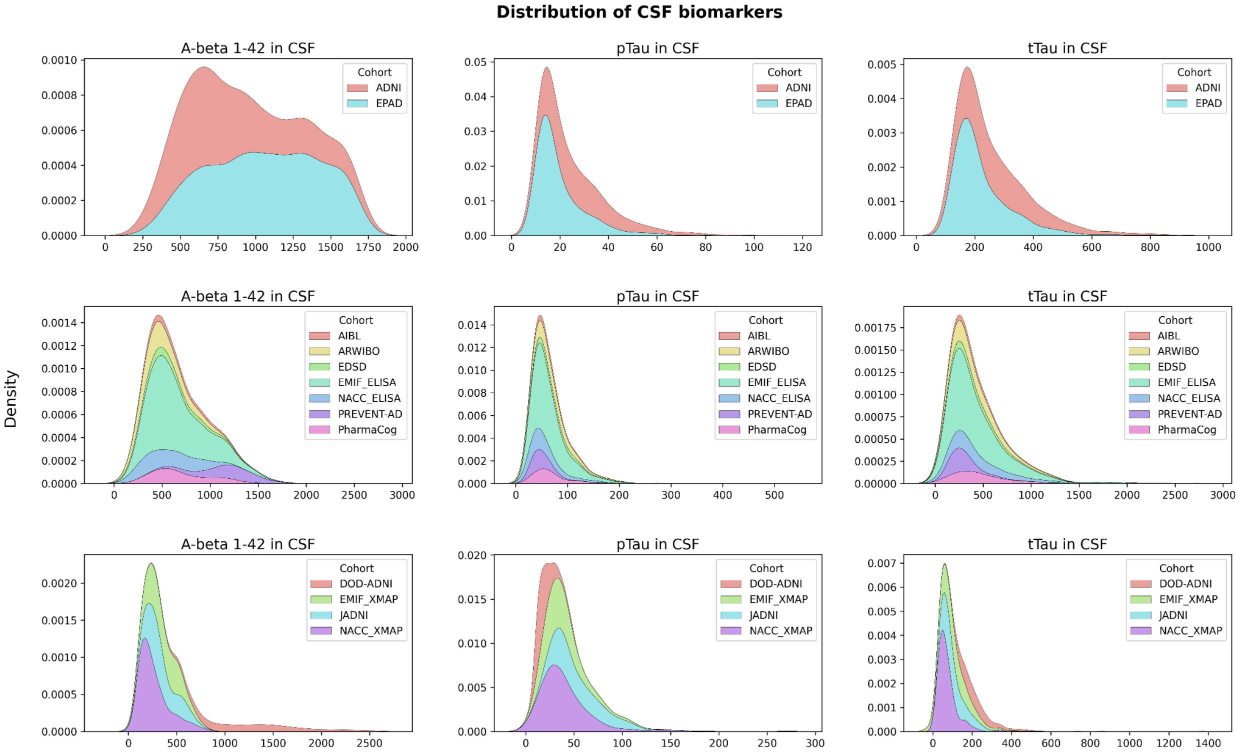


**Figure S1:** The distribution of CSF biomarkers in cohort studies. Note: the cohorts with the same assay method are grouped together in order to compare their distribution.

| Method | Number of Publication | Citation |
|---|---|---|
| **ROC using Youden's index** | 15 | [12-26] |
| **ROC** | 2 | [27, 28] |
| **ROC using Youden's index and mean ±2 SD** | 1 | [29] |
| **GMM, Youden's index and mean ±2 SD** | 1 | [30] |
| **Youden's index** | 2 | [31-32] |
| **GMM and mean ± SD** | 1 | [33] |
| **GMM, ROC using Youden's index** | 1 | [34] |
| **GMM and ROC** | 1 | [35] |
| **Sparse K-means** | 1 | [36] |
| **ROC using Youden's index and regression analysis** | 1 | [37] |
| **Threshold from previous studies, GMM and ROC** | 1 | [38] |
| **Tertile and quartiles** | 1 | [39] |
| **Threshold from previous studies** | 26 | [40-65] |
| **Threshold from previous studies, mean ±2 SD and 90th percentile** | 1 | [66] |
| **Youden's index and mean ±2 SD** | 1 | [67] |

**Table S4:** Commonly used methodologies extracted from literature for defining thresholds of ATN biomarkers.

| **Common Features** | | | |
|---|---|---|---|
| Cortical White Matter Volume | Left Pars Triangularis Gray Matter Volume | Right Caudal Anterior Cingulate Mean Cortical Thickness | Right Parsorbitalis Mean Cortical Thickness |
| Left Caudal Anterior Cingulate Mean Cortical Thickness | Left Pars Triangularis Mean Cortical Thickness | Right Cuneus Mean Cortical Thickness | Right Pericalcarine Gray Matter Volume |
| Left Caudal Middle Frontal Mean Cortical Thickness | Left Parsorbitalis Mean Cortical Thickness | Right Entorightinal Mean Cortical Thickness | Right Pericalcarine Mean Cortical Thickness |
| Left Cuneus Mean Cortical Thickness | Left Pericalcarine Gray Matter Volume | Right Fusiform Mean Cortical Thickness | Right Postcentral Grey Matter Volume |

| | | | |
|---|---|---|---|
| Left Fusiform Mean Cortical Thickness | Left Pericalcarine Mean Cortical Thickness | Right Hippocampus Volume | Right Postcentral Mean Cortical Thickness |
| Left Hippocampus Volume | Left Postcentral Gray Matter Volume | Right Inferiorparietal Mean Cortical Thickness | Right Posterior Cingulate Gray Matter Volume |
| Left Inferiorparietal Mean Cortical Thickness | Left Postcentral Mean Cortical Thickness | Right Inferiortemporal Mean Cortical Thickness | Right Posterior Cingulate Mean Cortical Thickness |
| Left Inferiortemporal Mean Cortical Thickness | Left Posterior Cingulate Gray Matter Volume | Right Insula Mean Cortical Thickness | Right Precentral Gray Matter Volume |
| Left Insula Mean Cortical Thickness | Left Posterior Cingulate Mean Cortical Thickness | Right Isthmus Cingulate Mean Cortical Thickness | Right Precentral Mean Cortical Thickness |
| Left Isthmus Cingulate Mean Cortical Thickness | Left Precentral Gray Matter Volume | Right Lateral Occipital Gray Matter Volume | Right Precuneus Grey Matter Volume |
| Left Lateral Occipital Gray Matter Volume | Left Precentral Mean Cortical Thickness | Right Lateral Ventricle Volume | Right Precuneus Mean Cortical Thickness |
| Left Lateral Ventricle Volume | Left Precuneus Grey Matter Volume | Right Lateraloccipital Mean Cortical Thickness | Right Rostral Anterior Cingulate Gray Matter Volume |
| Left Lateraloccipital Mean Cortical Thickness | Left Precuneus Mean Cortical Thickness | Right Lateralorbitofrontal Mean Cortical Thickness | Right Rostral Anterior Cingulate Mean Cortical Thickness |
| Left Lateralorbitofrontal Mean Cortical Thickness | Left Rostral Anterior Cingulate Gray Matter Volume | Right Lingual Grey Matter Volume | Right Rostral Middle Frontal Gray Matter Volume |
| Left Lingual Grey Gray Matter Volume | Left Rostral Anterior Cingulate Mean Cortical Thickness | Right Lingual Mean Cortical Thickness | Right Rostral Middle Frontal Mean Cortical Thickness |
| Left Lingual Mean Cortical Thickness | Left Rostral Middle Frontal Gray Matter Volume | Right Medial Orbitofrontal Grey Matter Volume | Right Superior Frontal Gray Matter Volume |
| Left Medial Orbitofrontal Gray Matter Volume | Left Rostral Middle Frontal Mean Cortical Thickness | Right Medial Orbitofrontal Mean Cortical Thickness | Right Superior Parietal Gray Matter Volume |
| Left Medial Orbitofrontal Mean Cortical Thickness | Left Superior Frontal Gray Matter Volume | Right Middle Temporal Gray Matter Volume | Right Superior Temporal Gray Matter Volume |
| Left Middle Temporal Gray Matter Volume | Left Superior Frontal Mean Cortical Thickness | Right Middle Temporal Mean Cortical Thickness | Right Superior Temporal Mean Cortical Thickness |
| Left Middle Temporal Mean Cortical Thickness | Left Superior Parietal Gray Matter Volume | Right Paracentral Mean Cortical Thickness | Right Superiorfrontal Mean Cortical Thickness |

| | | | |
|---|---|---|---|
| Left Paracentral Gray Matter Volume | Left Superior Temporal Mean Cortical Thickness | Right Parahippocampal Mean Cortical Thickness | Right Superiorparietal Mean Cortical Thickness |
| Left Paracentral Mean Cortical Thickness | Left Superiorparietal Mean Cortical Thickness | Right Pars Opercularis Gray Matter Volume | Right Supramarginal Gray Matter Volume |
| Left Parahippocampal Mean Cortical Thickness | Left Supramarginal Gray Matter Volume | Right Pars Opercularis Mean Cortical Thickness | Right Supramarginal Mean Cortical Thickness |
| Left Pars Opercularis Gray Matter Volume | Left Supramarginal Mean Cortical Thickness | Right Pars Orbitalis Gray Matter Volume | Right Transverse Temporal Grey Matter Volume |
| Left Pars Opercularis Mean Cortical Thickness | Left Transverse Temporal Grey Matter Volume | Right Pars Triangularis Gray Matter Volume | Right Transverse Temporal Mean Cortical Thickness |
| Left Pars Orbitalis Gray Matter Volume | Left Transverse Temporal Mean Cortical Thickness | Right Pars Triangularis Mean Cortical Thickness | APOE ε4 allele status |
| Third Ventricle Volume | | | |

**Table S5:** The common attributes among the investigated cohorts that were used for clustering and UMAP analysis. Here, 104 MRI measurements and the APOE ε4 status of participants were utilized.

| Cohort | Method | | | | | | | | | | | | | | |
|---|---|---|---|---|---|---|---|---|---|---|---|---|---|---|---|
| | GMM | | | K-means | | | Tertile | | | ROC | | | Mean ±2 SD | | |
| | Aβ1-42 | pTau | tTau | Aβ1-42 | pTau | tTau | Aβ1-42 | pTau | tTau | Aβ1-42 | pTau | tTau | Aβ1-42 | pTau | tTau |
| **ADNI** | 986.9 [986.0, 987.9] | 34.7 [34.6, 34.8] | 356.9 [356.1, 357.7] | 972.6 [971.6, 973.6] | 28.1 [28.0, 28.1] | 288.2 [287.8, 288.6] | 833.3 [831.2, 835.3] | 21.8 [21.8, 21.9] | 240.1 [239.5, 240.6] | 819.4 [815.8, 823.1] | 23.6 [23.5, 23.7] | 259.0 [258.0, 260.0] | 283.8 [281.8, 285.8] | 40.7 [40.6, 40.8] | 408.5 [407.6, 409.4] |
| **EPAD** | 1035.7 [1035.0, 1036.4] | 27.8 [27.7, 27.8] | 307.0 [306.6, 307.4] | 1033.3 [1032.6, 1034.0] | 19.8 [19.8, 19.9] | 223.7 [223.5, 223.9] | 884.5 [883.5, 885.4] | 19.9 [19.8, 19.9] | 228.2 [227.9, 228.5] | 684.7 [680.5, 688.9] | 23.9 [23.6, 24.3] | 267.9 [263.3, 272.4] | 339.0 [338.1, 339.9] | 43.3 [43.2, 43.4] | 435.9 [435.2, 436.7] |
| **AIBL** | 749.0 [743.2, 754.8] | 87.8 [86.8, 88.9] | 604.5 [597.0, 612.0] | 684.0 [681.2, 686.7] | 77.7 [77.0, 78.4] | 530.6 [523.3, 537.9] | - | - | - | - | - | - | - | - | - |
| **ARWIBO** | 589.2 [585.0, 593.5] | 155.0 [148.2, 161.9] | 531.6 [526.7, 536.4] | 523.8 [521.7, 526.0] | 69.1 [68.9, 69.4] | 432.1 [428.8, 435.4] | - | - | - | - | - | - | - | - | - |
| **EDSD** | 782.2 [776.1, 788.3] | 119.2 [117.7, 120.8] | 584.8 [576.2, 593.5] | 741.8 [736.1, 747.5] | 86.7 [86.0, 87.4] | 547.6 [543.0, 552.2] | - | - | - | - | - | - | - | - | - |
| **PREVENT-AD** | 1187.4 [1158.6, 1216.2] | 71.1 [70.3, 72.0] | 466.5 [457.2, 475.8] | 1146.4 [1144.1, 1148.8] | 50.8 [50.6, 51.0] | 308.8 [306.6, 310.9] | 1098.7 [1096.8, 1100.6] | 53.0 [52.9, 53.1] | 303.3 [302.5, 304.1] | - | - | - | 593.6 [589.7, 597.6] | 89.2 [88.9, 89.6] | 593.3 [589.5, 597.2] |
| **PharmaCog** | 792.9 [791.1, 794.7] | 93.0 [92.7, 93.3] | 773.4 [750.8, 796.0] | 762.2 [759.5, 764.8] | 68.6 [68.0, 69.2] | 466.9 [464.6, 469.2] | - | - | - | - | - | - | - | - | - |
| **NACC_ELISA** | 594.1 [592.6, 595.7] | 81.7 [80.9, 82.5] | 564.6 [552.4, 576.9] | 586.1 [584.9, 587.2] | 61.6 [61.4, 61.8] | 492.1 [490.2, 493.9] | 625.5 [623.6, 627.5] | 48.3 [48.1, 48.4] | 371.3 [369.6, 373.0] | 451.5 [448.1, 454.8] | 61.2 [60.9, 61.6] | 500.6 [496.0, 505.1] | 294.4 [292.6, 296.2] | 87.3 [86.8, 87.8] | 725.3 [722.0, 728.7] |
| **EMIF_ELISA** | 742.1 [741.2, 743.0] | 82.1 [81.9, 82.3] | 526.6 [524.2, 528.9] | 697.7 [695.2, 700.1] | 64.9 [64.8, 65.0] | 432.5 [431.7, 433.4] | 541.0 [539.7, 542.2] | 52.3 [52.2, 52.4] | 268.8 [268.0, 269.6] | 580.2 [575.8, 584.5] | 62.6 [62.1, 63.1] | 335.7 [333.7, 337.8] | 175.2 [173.7, 176.8] | 94.8 [94.3, 95.3] | 653.8 [645.4, 662.1] |
| **NACC_xMAP** | 298.7 | 60.8 | 94.5 | 290.7 | 40.1 | 70.2 | 248.5 | 36.5 | 54.0 | 225.3 | 40.3 | 62.7 | 23.6 | 68.9 | 99.3 |

| | | | | | | | | | | | | | | | |
|---|---|---|---|---|---|---|---|---|---|---|---|---|---|---|---|
| | [297.0, 300.4] | [59.9, 61.7] | [93.8, 95.3] | [288.4, 293.0] | [40.0, 40.3] | [70.0, 70.4] | [247.7, 249.3] | [36.4, 36.6] | [53.8, 54.2] | [224.8, 225.9] | [40.1, 40.5] | [62.1, 63.3] | [22.5, 24.6] | [68.5, 69.3] | [98.8, 99.7] |
| **EMIF_xMAP** | 386.9 [384.2, 389.5] | 51.6 [51.2, 52.1] | 227.7 [220.8, 234.5] | 363.3 [362.3, 364.2] | 37.5 [37.4, 37.6] | 128.9 [128.3, 129.5] | 446.7 [444.9, 448.4] | 31.4 [31.1, 31.6] | 80.9 [80.3, 81.4] | 375.4 [373.2, 377.5] | 30.6 [30.3, 30.9] | 98.6 [98.0, 99.1] | 247.6 [244.6, 250.6] | 54.5 [54.1, 54.8] | 122.7 [122.1, 123.2] |
| **DOD-ADNI** | 1412.8 [1394.9, 1430.8] | 26.6 [26.4, 26.7] | 304.0 [302.3, 305.7] | 1273.5 [1267.1, 1279.8] | 22.4 [22.3, 22.5] | 249.8 [248.8, 250.8] | 928.4 [924.4, 932.4] | 19.8 [19.8, 19.9] | 232.7 [231.9, 233.4] | - | - | - | 289.3 [285.5, 293.2] | 34.2 [34.1, 34.4] | 369.6 [368.3, 370.8] |
| **JADNI** | 414.7 [412.5, 416.9] | 61.1 [60.8, 61.5] | 133.9 [132.6, 135.2] | 392.5 [390.6, 394.4] | 59.9 [59.6, 60.1] | 129.0 [127.9, 130.0] | 392.7 [390.9, 394.5] | 38.5 [38.5, 38.6] | 72.0 [71.8, 72.3] | 343.1 [341.7, 344.6] | 46.0 [45.8, 46.3] | 84.9 [84.7, 85.0] | 191.2 [189.5, 193.0] | 65.1 [64.5, 65.6] | 126.3 [125.4, 127.3] |

**Table S6:** Thresholds obtained using each methodology for CSF biomarkers through 1000 bootstraps. The mean thresholds and the confidence intervals, in brackets, are given.

| Cohort | Method | | | | | | | | | | | | | | |
|---|---|---|---|---|---|---|---|---|---|---|---|---|---|---|---|
| | GMM | | | K-means | | | Tertile | | | ROC | | | Mean ±2 SD | | |
| | Aβ1-42 | pTau | tTau | Aβ1-42 | pTau | tTau | Aβ1-42 | pTau | tTau | Aβ1-42 | pTau | tTau | Aβ1-42 | pTau | tTau |
| **ADNI** | 0.19 % | 0.64 % | 0.47 % | 0.21 % | 0.25 % | 0.26 % | 0.48 % | 0.5 % | 0.48 % | 0.89 % | 0.85 % | 0.77 % | 1.4 % | 0.53 % | 0.45 % |
| **EPAD** | 0.14 % | 0.33 % | 0.25 % | 0.13 % | 0.22 % | 0.2 % | 0.21 % | 0.28 % | 0.27 % | 1.23 % | 3.02 % | 3.39 % | 0.54 % | 0.43 % | 0.34 % |
| **AIBL** | 1.54 % | 2.39 % | 2.48 % | 0.8 % | 1.81 % | 2.75 % | - | - | - | - | - | - | - | - | - |
| **ARWIBO** | 1.44 % | 8.85 % | 1.84 % | 0.81 % | 0.72 % | 1.52 % | - | - | - | - | - | - | - | - | - |
| **EDSD** | 1.56 % | 2.59 % | 2.95 % | 1.54 % | 1.61 % | 1.68 % | - | - | - | - | - | - | - | - | - |
| **PREVENT-AD** | 4.84 % | 2.46 % | 3.99 % | 0.41 % | 0.9 % | 1.39 % | 0.35 % | 0.52 % | 0.51 % | - | - | - | 1.33 % | 0.87 % | 1.29 % |
| **PharmaCog** | 0.45 % | 0.61 % | 5.85 % | 0.69 % | 1.71 % | 0.98 % | - | - | - | - | - | - | - | - | - |
| **NACC_ELISA** | 0.51 % | 1.99 % | 4.33 % | 0.39 % | 0.69 % | 0.76 % | 0.62 % | 0.55 % | 0.9 % | 1.47 % | 1.16 % | 1.82 % | 1.22 % | 1.11 % | 0.93 % |
| **EMIF_ELISA** | 0.24 % | 0.46 % | 0.9 % | 0.7 % | 0.27 % | 0.38 % | 0.45 % | 0.39 % | 0.57 % | 1.51 % | 1.51 % | 1.22 % | 1.78 % | 1.08 % | 2.56 % |
| **NACC_xMAP** | 1.15 % | 2.99 % | 1.54 % | 1.56 % | 0.64 % | 0.64 % | 0.63 % | 0.38 % | 0.61 % | 0.48 % | 0.92 % | 1.8 % | 8.67 % | 1.19 % | 0.97 % |
| **EMIF_xMAP** | 1.36 % | 1.77 % | 6.02 % | 0.52 % | 0.53 % | 0.87 % | 0.77 % | 1.54 % | 1.37 % | 1.14 % | 1.71 % | 1.14 % | 2.45 % | 1.38 % | 0.84 % |
| **DOD-ADNI** | 2.55 % | 0.9 % | 1.13 % | 0.99 % | 0.97 % | 0.81 % | 0.85 % | 0.65 % | 0.62 % | - | - | - | 2.66 % | 0.84 % | 0.67 % |
| **JADNI** | 1.06 % | 1.05 % | 1.93 % | 0.96 % | 0.82 % | 1.66 % | 0.92 % | 0.45 % | 0.7 % | 0.85 % | 1.09 % | 0.41 % | 1.84 % | 1.58 % | 1.54 % |

**Table S7:** The variation among thresholds obtained through bootstrapping for each biomarker using different methodologies.

| Cohort | Biomarkers | | |
|---|---|---|---|
| | Aβ1-42 | pTau | tTau |
| **ADNI** | 71.45 | 47.43 | 41.3 |
| **EPAD** | 67.35 | 54.27 | 51.01 |
| **PREVENT-AD** | 48.85 | 43.86 | 49.57 |
| **NACC_ELISA** | 53.66 | 45.45 | 48.33 |
| **EMIF_ELISA** | 76.32 | 45.22 | 59.76 |
| **NACC_xMAP** | 92.7 | 47.62 | 46.04 |
| **EMIF_xMAP** | 46.8 | 48.01 | 59.26 |
| **DOD-ADNI** | 77.87 | 42.27 | 38.09 |
| **JADNI** | 55.12 | 41.24 | 50.48 |
| **Average** | 65.57 | 46.15 | 49.32 |

**Table S8:** The relative change among thresholds achieved through different methods for each biomarker. Percentages are calculated with respect to each cohort's largest threshold for the respective biomarker. AIBL, ARWIBO, EDSD, and PharmaCog were removed from this assessment as only two out of the five thresholding methods could be performed on them.

| Groups | Biomarkers | | |
|---|---|---|---|
| | Aβ1-42 | pTau | tTau |
| **ADNI**<br>**EPAD** | 8.39 % | 15.08 % | 13.05 % |
| **AIBL**<br>**ARWIBO**<br>**EDSD**<br>**PharmaCog**<br>**PREVENT-AD**<br>**NACC_ELISA**<br>**EMIF_ELISA** | 47.32 % | 22.05 % | 31.13 % |
| **NACC_xMAP**<br>**EMIF_xMAP**<br>**DOD-ADNI**<br>**JADNI** | 71.25 % | 51.23 % | 66.85 % |

**Table S9:** The relative changes of obtained thresholds across the cohort with the same employed assay for each biomarker.

| Groups | Method | | | | |
|---|---|---|---|---|---|
| | GMM | K-means | Tertile | ROC | Mean ±2 SD |

| | | | | | |
|---|---|---|---|---|---|
| **ADNI**<br>**EPAD** | 12.38 % | 18.99 % | 6.09 % | 13.89 % | 9.52 % |
| **AIBL**<br>**ARWIBO**<br>**EDSD**<br>**PharmaCog**<br>**PREVENT-AD**<br>**NACC_ELISA**<br>**EMIF_ELISA** | 40.91 % | 49.07 % | 29.78 % | 15.74 % | 31.99 % |
| **NACC_xMAP**<br>**EMIF_xMAP**<br>**DOD-ADNI**<br>**JADNI** | 67.55 % | 71.26 % | 66.4 % | 38.39 % | 71.95 % |

**Table S10:** The relative changes of obtained thresholds across the cohort with the same employed assay within each method.

| Cohort | ATN Biomarker Profile | | | | | | | |
|---|---|---|---|---|---|---|---|---|
| | A-T-N- | A-T+N+ | A-T-N+ | A-T+N- | A+T+N- | A+T-N- | A+T-N+ | A+T+N+ |
| ADNI | 275 | 28 | 5 | 2 | 25 | 443 | 12 | 227 |
| EPAD | 590 | 53 | 4 | 6 | 19 | 413 | 6 | 146 |
| AIBL | 17 | 0 | 2 | 0 | 0 | 28 | 1 | 9 |
| ARWIBO | 55 | 2 | 13 | 1 | 1 | 87 | 46 | 12 |
| EDSD | 21 | 1 | 4 | 1 | 2 | 36 | 9 | 11 |
| PREVENT-AD | 86 | 9 | 0 | 3 | 0 | 29 | 1 | 5 |
| PharmaCog | 46 | 1 | 2 | 0 | 1 | 67 | 5 | 23 |
| NACC | 168 | 6 | 9 | 3 | 12 | 193 | 51 | 64 |
| EMIF | 298 | 18 | 11 | 8 | 40 | 417 | 61 | 161 |
| DOD-ADNI | 46 | 8 | 0 | 1 | 0 | 50 | 0 | 8 |
| JADNI | 55 | 0 | 1 | 0 | 20 | 65 | 5 | 51 |

**Table S11:** The number of categorized participants in each ATN profile using thresholds that were obtained by GMM methodology.

| Cohort | ATN Biomarker Profile | | | | | | | |
|---|---|---|---|---|---|---|---|---|
| | A-T-N- | A-T+N+ | A-T-N+ | A-T+N- | A+T+N- | A+T-N- | A+T-N+ | A+T+N+ |
| ADNI | 256 | 49 | 14 | 0 | 18 | 305 | 16 | 359 |
| EPAD | 503 | 121 | 28 | 2 | 22 | 277 | 7 | 277 |
| AIBL | 17 | 1 | 1 | 3 | 1 | 23 | 1 | 10 |

| | | | | | | | | |
|---|---|---|---|---|---|---|---|---|
| **ARWIBO** | 55 | 18 | 5 | 7 | 8 | 50 | 15 | 59 |
| **EDSD** | 19 | 3 | 1 | 3 | 10 | 29 | 2 | 18 |
| **PREVENT-AD** | 39 | 30 | 3 | 6 | 2 | 39 | 1 | 13 |
| **PharmaCog** | 40 | 5 | 4 | 0 | 9 | 41 | 3 | 43 |
| **NACC** | 144 | 23 | 13 | 24 | 28 | 123 | 47 | 104 |
| **EMIF** | 265 | 42 | 16 | 35 | 64 | 283 | 37 | 272 |
| **DOD-ADNI** | 36 | 18 | 4 | 0 | 1 | 42 | 3 | 9 |
| **JADNI** | 56 | 0 | 1 | 0 | 29 | 67 | 2 | 42 |

**Table S12:** The number of categorized participants in each ATN profile using thresholds that were obtained by the K-means methodology.

| **Cohort** | **ATN Biomarker Profile** | | | | | | | |
|---|---|---|---|---|---|---|---|---|
| | **A-T-N-** | **A-T+N+** | **A-T-N+** | **A-T+N-** | **A+T+N-** | **A+T-N-** | **A+T-N+** | **A+T+N+** |
| **ADNI** | 265 | 160 | 10 | 3 | 33 | 148 | 3 | 395 |
| **EPAD** | 621 | 167 | 24 | 6 | 24 | 169 | 5 | 221 |
| **PREVENT-AD** | 52 | 29 | 3 | 5 | 1 | 31 | 2 | 10 |
| **NACC** | 109 | 56 | 19 | 25 | 10 | 88 | 53 | 146 |
| **EMIF** | 207 | 220 | 63 | 29 | 13 | 118 | 46 | 318 |
| **DOD-ADNI** | 46 | 27 | 1 | 1 | 3 | 24 | 2 | 9 |
| **JADNI** | 32 | 19 | 9 | 2 | 11 | 14 | 8 | 102 |

**Table S13:** The number of categorized participants in each ATN profile using thresholds that were obtained by tertile methodology.

| **Cohort** | **ATN Biomarker Profile** | | | | | | | |
|---|---|---|---|---|---|---|---|---|
| | **A-T-N-** | **A-T+N+** | **A-T-N+** | **A-T+N-** | **A+T+N-** | **A+T-N-** | **A+T-N+** | **A+T+N+** |
| **ADNI** | 304 | 147 | 8 | 6 | 34 | 180 | 1 | 337 |
| **EPAD** | 639 | 223 | 85 | 0 | 3 | 120 | 13 | 154 |
| **NACC** | 182 | 40 | 32 | 11 | 7 | 65 | 67 | 102 |
| **EMIF** | 294 | 148 | 39 | 53 | 22 | 148 | 27 | 283 |
| **JADNI** | 61 | 19 | 9 | 0 | 10 | 16 | 5 | 77 |

**Table S14:** The number of categorized participants in each ATN profile using thresholds that were obtained by ROC (Youdnen's index) methodology.

| Cohort | ATN Biomarker Profile | | | | | | | |
|---|---|---|---|---|---|---|---|---|
| | A-T-N- | A-T+N+ | A-T-N+ | A-T+N- | A+T+N- | A+T-N- | A+T-N+ | A+T+N+ |
| ADNI | 813 | 154 | 20 | 22 | 0 | 7 | 0 | 1 |
| EPAD | 1165 | 49 | 9 | 4 | 0 | 10 | 0 | 0 |
| PREVENT-AD | 123 | 5 | 1 | 1 | 0 | 3 | 0 | 0 |
| NACC | 385 | 35 | 43 | 13 | 2 | 17 | 2 | 9 |
| EMIF | 742 | 117 | 70 | 33 | 1 | 31 | 15 | 5 |
| DOD-ADNI | 106 | 5 | 1 | 1 | 0 | 0 | 0 | 0 |
| JADNI | 119 | 48 | 8 | 4 | 3 | 10 | 1 | 4 |

**Table S15:** The number of categorized participants in each ATN profile using thresholds that were obtained by mean ±2 SD methodology.

| Cohorts | GMM | | | | | | | |
|---|---|---|---|---|---|---|---|---|
| | A-T-N- | A-T+N+ | A-T-N+ | A+T+N- | A+T-N- | A+T-N+ | A+T+N+ | Total |
| ADNI | 170 | 18 | 2 | 13 | 256 | 8 | 122 | 589 |
| EDSD | 14 | 0 | 3 | 1 | 27 | 8 | 10 | 63 |
| ARWIBO | 23 | 2 | 7 | 0 | 27 | 14 | 7 | 80 |
| NACC | 80 | 2 | 7 | 1 | 62 | 27 | 17 | 196 |
| JADNI | 52 | 0 | 1 | 17 | 63 | 5 | 49 | 187 |
| DOD-ADNI | 28 | 4 | 0 | 0 | 34 | 0 | 4 | 70 |
| PharmaCog | 46 | 1 | 2 | 1 | 65 | 5 | 23 | 143 |
| Total | 413 | 27 | 22 | 33 | 534 | 67 | 232 | 1328 |

**Table S16:** The number of participants included in the ATN-based analysis in each cohort using GMM thresholds.

| Cohorts | K-means | | | | | | | |
|---|---|---|---|---|---|---|---|---|
| | A-T-N- | A-T+N+ | A-T-N+ | A+T+N- | A+T-N- | A+T-N+ | A+T+N+ | Total |
| ADNI | 161 | 30 | 9 | 9 | 181 | 7 | 194 | 591 |
| EDSD | 12 | 2 | 0 | 8 | 22 | 2 | 16 | 62 |

| | | | | | | | | |
|---|---|---|---|---|---|---|---|---|
| **ARWIBO** | 18 | 8 | 3 | 3 | 17 | 4 | 22 | 75 |
| **NACC** | 71 | 9 | 2 | 3 | 48 | 21 | 34 | 188 |
| **JADNI** | 53 | 0 | 1 | 26 | 65 | 2 | 40 | 187 |
| **DOD-ADNI** | 23 | 7 | 3 | 1 | 31 | 0 | 5 | 70 |
| **PharmaCog** | 40 | 5 | 4 | 9 | 40 | 3 | 42 | 143 |
| **Total** | 378 | 61 | 22 | 59 | 404 | 39 | 353 | 1316 |

**Table S17:** The number of participants included in the ATN-based analysis in each cohort using K-means thresholds.

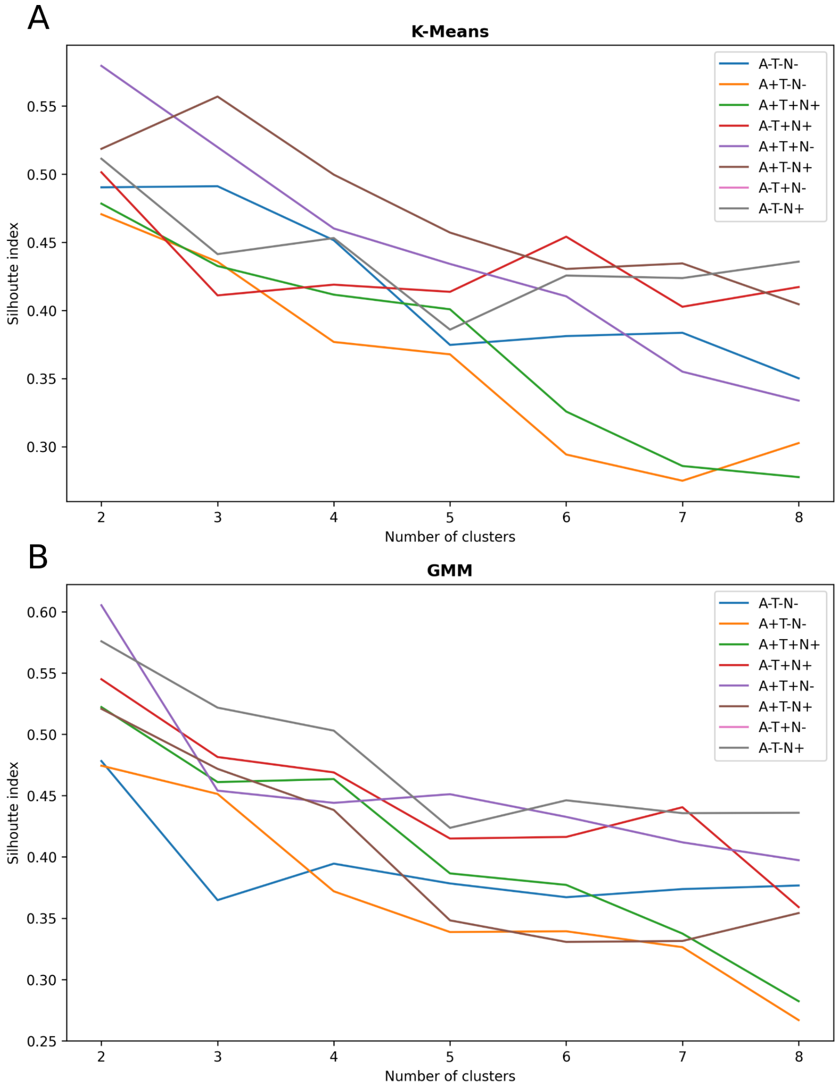


**Figure S2:** Silhouette index for different numbers of clusters within each ATN profile using GMM and K-means thresholds. A) Using K-means thresholds for the categorization of participants. B) Using GMM thresholds for the categorization of participants.

| Cohort | Cluster (A-T-N-) | | |
|---|---|---|---|
| | 1 | 2 | 3 |
| ADNI | 50 | 85 | 26 |
| EDSD | 8 | 3 | 1 |
| ARWIBO | 6 | 9 | 3 |
| NACC | 24 | 35 | 12 |
| JADNI | 18 | 24 | 11 |
| DOD-ADNI | 8 | 12 | 3 |
| PharmaCog | 17 | 17 | 6 |

**Table S18:** The number of participants clustered with each cluster label (using silhouette score) within the A-T-N- group using K-means thresholds.

| Cohort | Cluster (A+T-N-) | |
|---|---|---|
| | 1 | 2 |
| ADNI | 105 | 76 |
| EDSD | 15 | 7 |
| ARWIBO | 13 | 4 |
| NACC | 28 | 20 |
| JADNI | 40 | 25 |
| DOD-ADNI | 22 | 9 |
| PharmaCog | 25 | 15 |

**Table S19:** The number of participants clustered with each cluster label (using silhouette score) within the A+T-N- group using K-means thresholds.

| Cohort | Cluster (A+T+N+) | |
|---|---|---|
| | 1 | 2 |
| ADNI | 108 | 86 |
| EDSD | 8 | 8 |
| ARWIBO | 14 | 8 |
| NACC | 18 | 16 |

| | | |
|---|---|---|
| **JADNI** | 24 | 16 |
| **DOD-ADNI** | 3 | 2 |
| **PharmaCog** | 23 | 19 |

**Table S20:** The number of participants clustered with each cluster label (using silhouette score) within the A+T+N+ group using K-means thresholds.

| **Cohort** | **Cluster (A-T+N+)** | |
|---|---|---|
| | **1** | **2** |
| **ADNI** | 22 | 8 |
| **EDSD** | 2 | 0 |
| **ARWIBO** | 6 | 2 |
| **NACC** | 6 | 3 |
| **DOD-ADNI** | 3 | 4 |
| **PharmaCog** | 2 | 3 |

**Table S21:** The number of participants clustered with each cluster label (using silhouette score) within the A-T+N+ group using K-means thresholds.

| **Cohort** | **Cluster (A+T+N-)** | |
|---|---|---|
| | **1** | **2** |
| **ADNI** | 2 | 7 |
| **EDSD** | 2 | 6 |
| **ARWIBO** | 0 | 3 |
| **NACC** | 1 | 2 |
| **JADNI** | 1 | 25 |
| **DOD-ADNI** | 0 | 1 |
| **PharmaCog** | 4 | 5 |

**Table S22:** The number of participants clustered with each cluster label (using silhouette score) within the A+T+N- group using K-means thresholds.

| **Cohort** | **Cluster (A-T-N+)** | |
|---|---|---|
| | **1** | **2** |

| | | |
|---|---|---|
| ADNI | 8 | 1 |
| ARWIBO | 1 | 2 |
| NACC | 2 | 0 |
| JADNI | 1 | 0 |
| DOD-ADNI | 1 | 2 |
| PharmaCog | 3 | 1 |

**Table S23:** The number of participants clustered with each cluster label (using silhouette score) within the A-T-N+ group using K-means thresholds.

| Cohort | Cluster (A+T-N+) | | |
|---|---|---|---|
| | 1 | 2 | 3 |
| ADNI | 3 | 2 | 2 |
| EDSD | 0 | 2 | 0 |
| ARWIBO | 3 | 0 | 1 |
| NACC | 7 | 8 | 6 |
| JADNI | 1 | 1 | 0 |
| PharmaCog | 1 | 1 | 1 |

**Table S24:** The number of participants clustered with each cluster label (using silhouette score) within the A+T-N+ group using K-means thresholds.

| Cohort | Cluster (A-T-N-) | |
|---|---|---|
| | 1 | 2 |
| ADNI | 106 | 64 |
| EDSD | 13 | 1 |
| ARWIBO | 15 | 8 |
| NACC | 47 | 33 |
| JADNI | 29 | 23 |
| DOD-ADNI | 18 | 10 |
| PharmaCog | 35 | 11 |

**Table S25:** The number of participants clustered with each cluster label (using silhouette score) within the A-T-N- group using GMM thresholds.

| Cohort | Cluster (A+T-N-) | |
|---|---|---|
| | 1 | 2 |
| ADNI | 156 | 100 |
| EDSD | 18 | 9 |
| ARWIBO | 19 | 8 |
| NACC | 38 | 24 |
| JADNI | 40 | 23 |
| DOD-ADNI | 23 | 11 |
| PharmaCog | 44 | 21 |

**Table S26:** The number of participants clustered with each cluster label (using silhouette score) within the A+T-N- group using GMM thresholds.

| Cohort | Cluster (A+T+N+) | |
|---|---|---|
| | 1 | 2 |
| ADNI | 95 | 27 |
| EDSD | 6 | 4 |
| ARWIBO | 7 | 0 |
| NACC | 13 | 4 |
| JADNI | 39 | 10 |
| DOD-ADNI | 1 | 3 |
| PharmaCog | 21 | 2 |

**Table S27:** The number of participants clustered with each cluster label (using silhouette score) within the A+T+N+ group using GMM thresholds.

| Cohort | Cluster (A-T+N+) | |
|---|---|---|
| | 1 | 2 |
| ADNI | 14 | 4 |
| ARWIBO | 2 | 0 |
| NACC | 2 | 0 |

| | | |
|---|---|---|
| **DOD-ADNI** | 3 | 1 |
| **PharmaCog** | 1 | 0 |

**Table S28:** The number of participants clustered with each cluster label (using silhouette score) within the A-T+N+ group using GMM thresholds.

| **Cohort** | **Cluster (A+T+N-)** | |
|---|---|---|
| | **1** | **2** |
| **ADNI** | 11 | 2 |
| **EDSD** | 0 | 1 |
| **NACC** | 1 | 0 |
| **JADNI** | 17 | 0 |
| **PharmaCog** | 0 | 1 |

**Table S29:** The number of participants clustered with each cluster label (using silhouette score) within the A+T+N- group using GMM thresholds.

| **Cohort** | **Cluster (A-T-N+)** | |
|---|---|---|
| | **1** | **2** |
| **ADNI** | 2 | 0 |
| **EDSD** | 2 | 1 |
| **ARWIBO** | 3 | 4 |
| **NACC** | 2 | 5 |
| **JADNI** | 0 | 1 |
| **PharmaCog** | 0 | 2 |

**Table S30:** The number of participants clustered with each cluster label (using silhouette score) within the A-T-N+ group using GMM thresholds.

| **Cohort** | **Cluster (A+T-N+)** | |
|---|---|---|
| | **1** | **2** |
| **ADNI** | 5 | 3 |
| **EDSD** | 7 | 1 |
| **ARWIBO** | 6 | 8 |

| | | |
|---|---|---|
| **NACC** | 18 | 9 |
| **JADNI** | 3 | 2 |
| **PharmaCog** | 2 | 3 |

**Table S31:** The number of participants clustered with each cluster label (using silhouette score) within the A+T-N+ group using GMM thresholds.

| **Method** | | **Biomarker Profile** | | | | | | |
|---|---|---|---|---|---|---|---|---|
| | | **A-T-N-** | **A-T+N+** | **A-T-N+** | **A+T+N-** | **A+T-N-** | **A+T-N+** | **A+T+N+** |
| **K-means** | **Cramer's V** | 0.09 | 0.13 | 0.21 | 0.16 | 0.1 | 0.07 | 0.11 |
| | ***p*-value** | 0.33 | 0.69 | 0.2 | 0.2 | 0.1 | 0.62 | 0.35 |
| | **Total Participants** | 378 | 61 | 22 | 59 | 404 | 39 | 353 |
| **GMM** | **Cramer's V** | 0.1 | 0.21 | 0.06 | 0.28 | 0.08 | 0.16 | 0.15 |
| | ***p*-value** | 0.1 | 0.69 | 0.41 | 0.08 | 0.3 | 0.12 | 0.05 |
| | **Total Participants** | 413 | 27 | 22 | 33 | 534 | 67 | 232 |

**Table S32:** The Cramer's V and the p-value for each clustering of participants in each ATN profile using certain data-driven thresholds. Note: Here, the number of clusters equals the number of cohorts within that profile.

| **Cohort** | **Cluster (A-T-N-)** | | | | | | |
|---|---|---|---|---|---|---|---|
| | **1** | **2** | **3** | **4** | **5** | **6** | **7** |
| **ADNI** | 35 | 15 | 46 | 1 | 39 | 13 | 12 |
| **EDSD** | 8 | 0 | 0 | 0 | 3 | 0 | 1 |
| **ARWIBO** | 4 | 2 | 5 | 0 | 4 | 3 | 0 |
| **NACC** | 20 | 4 | 23 | 1 | 12 | 8 | 3 |
| **JADNI** | 15 | 3 | 13 | 0 | 11 | 10 | 1 |
| **DOD-ADNI** | 4 | 4 | 6 | 1 | 6 | 2 | 0 |
| **PharmaCog** | 14 | 3 | 9 | 0 | 8 | 3 | 3 |

**Table S33:** The number of participants clustered with each cluster label within the A-T-N- group using K-means thresholds.

| **Cohort** | **Cluster (A+T-N-)** | | | | | | |
|---|---|---|---|---|---|---|---|
| | **1** | **2** | **3** | **4** | **5** | **6** | **7** |

| | | | | | | | |
|---|---|---|---|---|---|---|---|
| **ADNI** | 35 | 37 | 16 | 39 | 6 | 35 | 13 |
| **EDSD** | 2 | 1 | 5 | 6 | 2 | 6 | 0 |
| **ARWIBO** | 2 | 1 | 4 | 3 | 1 | 4 | 2 |
| **NACC** | 14 | 10 | 3 | 10 | 1 | 3 | 7 |
| **JADNI** | 17 | 11 | 4 | 14 | 2 | 7 | 10 |
| **DOD-ADNI** | 7 | 4 | 2 | 5 | 0 | 11 | 2 |
| **PharmaCog** | 11 | 5 | 5 | 10 | 0 | 6 | 3 |

**Table S34:** The number of participants clustered with each cluster label within the A+T-N- group using K-means thresholds.

| **Cohort** | **Cluster (A+T+N+)** | | | | | | |
|---|---|---|---|---|---|---|---|
| | **1** | **2** | **3** | **4** | **5** | **6** | **7** |
| **ADNI** | 29 | 34 | 13 | 35 | 34 | 26 | 23 |
| **EDSD** | 0 | 1 | 4 | 2 | 4 | 1 | 4 |
| **ARWIBO** | 2 | 6 | 3 | 1 | 1 | 4 | 5 |
| **NACC** | 6 | 6 | 1 | 3 | 6 | 8 | 4 |
| **JADNI** | 5 | 6 | 1 | 11 | 5 | 6 | 6 |
| **DOD-ADNI** | 1 | 2 | 0 | 0 | 0 | 1 | 1 |
| **PharmaCog** | 6 | 5 | 2 | 10 | 6 | 6 | 7 |

**Table S35:** The number of participants clustered with each cluster label within the A+T+N+ group using K-means thresholds.

| **Cohort** | **Cluster (A-T+N+)** | | | | | |
|---|---|---|---|---|---|---|
| | **1** | **2** | **3** | **4** | **5** | **6** |
| **ADNI** | 8 | 6 | 10 | 2 | 3 | 1 |
| **EDSD** | 1 | 0 | 1 | 0 | 0 | 0 |
| **ARWIBO** | 4 | 2 | 1 | 0 | 1 | 0 |
| **NACC** | 4 | 3 | 1 | 0 | 1 | 0 |
| **DOD-ADNI** | 1 | 3 | 1 | 1 | 0 | 1 |
| **PharmaCog** | 0 | 3 | 0 | 0 | 1 | 1 |

**Table S36:** The number of participants clustered with each cluster label within the A-T+N+ group using K-means thresholds.

| Cohort | Cluster (A+T+N-) | | | | | | |
|---|---|---|---|---|---|---|---|
| | 1 | 2 | 3 | 4 | 5 | 6 | 7 |
| **ADNI** | 4 | 0 | 0 | 1 | 2 | 0 | 2 |
| **EDSD** | 1 | 2 | 0 | 2 | 3 | 0 | 0 |
| **ARWIBO** | 2 | 0 | 1 | 0 | 0 | 0 | 0 |
| **NACC** | 1 | 0 | 0 | 0 | 1 | 0 | 1 |
| **JADNI** | 6 | 0 | 7 | 5 | 7 | 1 | 0 |
| **DOD-ADNI** | 0 | 0 | 0 | 1 | 0 | 0 | 0 |
| **PharmaCog** | 2 | 3 | 0 | 1 | 2 | 0 | 1 |

**Table S37:** The number of participants clustered with each cluster label within the A+T+N- group using K-means thresholds.

| Cohort | Cluster (A-T-N+) | | | | | |
|---|---|---|---|---|---|---|
| | 1 | 2 | 3 | 4 | 5 | 6 |
| **ADNI** | 3 | 4 | 0 | 0 | 1 | 1 |
| **ARWIBO** | 0 | 0 | 1 | 0 | 1 | 1 |
| **NACC** | 1 | 0 | 0 | 0 | 1 | 0 |
| **JADNI** | 0 | 1 | 0 | 0 | 0 | 0 |
| **DOD-ADNI** | 0 | 1 | 2 | 0 | 0 | 0 |
| **PharmaCog** | 3 | 0 | 0 | 1 | 0 | 0 |

**Table S38:** The number of participants clustered with each cluster label within the A-T-N+ group using K-means thresholds.

| Cohort | Cluster (A+T-N+) | | | | | |
|---|---|---|---|---|---|---|
| | 1 | 2 | 3 | 4 | 5 | 6 |
| **ADNI** | 2 | 2 | 2 | 1 | 0 | 0 |
| **EDSD** | 0 | 2 | 0 | 0 | 0 | 0 |
| **ARWIBO** | 0 | 0 | 1 | 1 | 0 | 2 |

| | | | | | | |
|---|---|---|---|---|---|---|
| **NACC** | 4 | 5 | 6 | 1 | 3 | 2 |
| **JADNI** | 0 | 1 | 0 | 0 | 0 | 1 |
| **PharmaCog** | 1 | 1 | 1 | 0 | 0 | 0 |

**Table S39:** The number of participants clustered with each cluster label within the A+T-N+ group using K-means thresholds.

| Cohort | Cluster (A-T-N-) | | | | | | |
|---|---|---|---|---|---|---|---|
| | 1 | 2 | 3 | 4 | 5 | 6 | 7 |
| **ADNI** | 12 | 19 | 52 | 17 | 33 | 35 | 2 |
| **EDSD** | 1 | 0 | 4 | 0 | 9 | 0 | 0 |
| **ARWIBO** | 0 | 6 | 7 | 3 | 2 | 5 | 0 |
| **NACC** | 4 | 10 | 20 | 9 | 17 | 20 | 0 |
| **JADNI** | 1 | 5 | 11 | 8 | 13 | 14 | 0 |
| **DOD-ADNI** | 1 | 7 | 7 | 2 | 3 | 7 | 1 |
| **PharmaCog** | 3 | 5 | 16 | 2 | 13 | 6 | 1 |

**Table S40:** The number of participants clustered with each cluster label within the A-T-N- group using GMM thresholds.

| Cohort | Cluster (A+T-N-) | | | | | | |
|---|---|---|---|---|---|---|---|
| | 1 | 2 | 3 | 4 | 5 | 6 | 7 |
| **ADNI** | 77 | 43 | 28 | 40 | 44 | 8 | 16 |
| **EDSD** | 8 | 2 | 6 | 1 | 8 | 2 | 0 |
| **ARWIBO** | 7 | 4 | 7 | 3 | 4 | 1 | 1 |
| **NACC** | 18 | 13 | 6 | 8 | 10 | 1 | 6 |
| **JADNI** | 16 | 18 | 4 | 10 | 12 | 2 | 1 |
| **DOD-ADNI** | 17 | 2 | 4 | 3 | 6 | 0 | 2 |
| **PharmaCog** | 23 | 12 | 8 | 7 | 11 | 1 | 3 |

**Table S41:** The number of participants clustered with each cluster label within the A+T-N- group using GMM thresholds.

| Cohort | Cluster (A+T+N+) | | | | | | |
|---|---|---|---|---|---|---|---|
| | 1 | 2 | 3 | 4 | 5 | 6 | 7 |
| ADNI | 43 | 11 | 8 | 0 | 30 | 19 | 11 |
| EDSD | 1 | 0 | 3 | 0 | 5 | 1 | 0 |
| ARWIBO | 1 | 2 | 0 | 0 | 2 | 0 | 2 |
| NACC | 6 | 3 | 0 | 0 | 4 | 4 | 0 |
| JADNI | 19 | 5 | 2 | 1 | 9 | 7 | 6 |
| DOD-ADNI | 0 | 1 | 0 | 0 | 0 | 3 | 0 |
| PharmaCog | 9 | 3 | 1 | 0 | 8 | 1 | 1 |

**Table S42:** The number of participants clustered with each cluster label within the A+T+N+ group using GMM thresholds.

| Cohort | Cluster (A-T+N+) | | | | |
|---|---|---|---|---|---|
| | 1 | 2 | 3 | 4 | 5 |
| ADNI | 5 | 4 | 7 | 0 | 2 |
| ARWIBO | 0 | 0 | 1 | 0 | 1 |
| NACC | 1 | 0 | 1 | 0 | 0 |
| DOD-ADNI | 1 | 0 | 1 | 1 | 1 |
| PharmaCog | 0 | 0 | 1 | 0 | 0 |

**Table S43:** The number of participants clustered with each cluster label within the A-T+N+ group using GMM thresholds.

| Cohort | Cluster (A+T+N-) | | | | |
|---|---|---|---|---|---|
| | 1 | 2 | 3 | 4 | 5 |
| ADNI | 7 | 2 | 1 | 2 | 1 |
| EDSD | 0 | 1 | 0 | 0 | 0 |
| NACC | 0 | 0 | 0 | 1 | 0 |
| JADNI | 7 | 0 | 6 | 3 | 1 |
| PharmaCog | 0 | 1 | 0 | 0 | 0 |

**Table S44:** The number of participants clustered with each cluster label within the A+T+N- group using GMM thresholds.

| Cohort | Cluster (A-T-N+) | | | | | |
|---|---|---|---|---|---|---|
| | 1 | 2 | 3 | 4 | 5 | 6 |
| ADNI | 0 | 0 | 2 | 0 | 0 | 0 |
| EDSD | 0 | 1 | 0 | 1 | 0 | 1 |
| ARWIBO | 0 | 2 | 2 | 1 | 2 | 0 |
| NACC | 2 | 2 | 1 | 0 | 1 | 1 |
| JADNI | 1 | 0 | 0 | 0 | 0 | 0 |
| PharmaCog | 0 | 1 | 0 | 0 | 1 | 0 |

**Table S45:** The number of participants clustered with each cluster label within the A-T-N+ group using GMM thresholds.

| Cohort | Cluster (A+T-N+) | | | | | |
|---|---|---|---|---|---|---|
| | 1 | 2 | 3 | 4 | 5 | 6 |
| ADNI | 3 | 2 | 0 | 1 | 0 | 2 |
| EDSD | 2 | 4 | 0 | 1 | 1 | 0 |
| ARWIBO | 2 | 4 | 2 | 3 | 0 | 3 |
| NACC | 9 | 4 | 6 | 2 | 5 | 1 |
| JADNI | 0 | 3 | 1 | 0 | 0 | 1 |
| PharmaCog | 0 | 1 | 0 | 0 | 1 | 3 |

**Table S46:** The number of participants clustered with each cluster label within the A+T-N+ group using GMM thresholds.
